# Baseline Inventory of the Bryophytes of Grand Staircase-Escalante National Monument, Utah

**DOI:** 10.64898/2026.02.27.708354

**Authors:** Theresa A. Clark

**Affiliations:** School of Life Sciences, University of Nevada, Las Vegas, NV 89154-4004

## Abstract

This report presents a preliminary bryoflora for Grand Staircase-Escalante National Monument (GSENM) in southern Utah. The inventory included over 1000 collections made across 40 localities (i.e. macrohabitat types) spanning two ecologically important gradients in bryophyte habitat: shade and moisture availability. At present, the growing checklist contains 117 taxa of liverworts and mosses including 27 families, 65 genera, 116 species, 9 varieties, and 1 subspecies. Noteworthy records include 49 putative taxa new for the state of Utah, and 2 undescribed species in the genera *Grimmia* and *Schistidium*. We propose 4 of these species be considered for addition to the recently revised bryoflora of North America. As expected for arid and semiarid environments, the bryophytes of GSENM are predominantly acrocarpous mosses (75%) followed by pleurocarpous mosses (16%), thalloid liverworts (7%), and leafy liverworts (2%). The most diverse families included xeric-soil acrocarpous mosses in the Pottiaceae (35%) and xeric-rock acrocarpous mosses in the Grimmiaceae (15%). Both xeric and mesic species were recovered in the Bryaceae (10% of species) while the pleurocarpous Amblystegiaceae included mesic and hydric species (7%). Most species in the bryoflora have broad global or disjunct distributions, but notably, the known distribution of 17 species appears limited in the United States, or globally, and warrant monitoring in GSENM.

Using floristic habitat sampling across 19 macrohabitat types (combinations of 6 topography and 7 vegetation classes), mean site richness was 17.2 ± 9 (SD) and ranged from 4 to 34 species. Six diversity hotspots supported ≥30 species and were canyons with perennial or ephemeral streams dominated by mixed conifer, hardwood-riparian, riparian, or pinyon-juniper vegetation. High richness is likely supported by greater habitat diversity including xeric, mesic, and hydric conditions on variable substrates (e.g. rock, soil, biocrust, downed wood, seeps, and riparian aquatic/semi-aquatic habitat). Consequently, managing and monitoring diversity under future climate change and land-use alterations will necessitate a habitat-stratified approach that utilizes repeated floristic habitat sampling to document changes in site-level richness and to predict other candidate diversity hotspots on the basis of microhabitat-level diversity, which could be assessed by trained non-bryologists.

Collection data are available to the public as georeferenced and photographed observations of half of the bryophyte collections on our iNaturalist.com project, *Bryophytes of Grand Staircase Escalante*, available for scientific, educational, or outreach activities. Observations are accessible to visitors (via the smartphone app) who wish to know what species have been found along popular trails in GSENM. Landscape-level richness may not reach that of the neighboring Grand Canyon National Park (>155 species), which supports a unique high-elevation bryophyte community sheltered in the mixed conifer and spruce-fir forests of the North Rim’s Kaibab Plateau. Future collecting by experts will inevitably uncover more species in this ecologically diverse monument important to conserving dryland bryophyte diversity and ecosystem function. This study will serve as a baseline for future research and long-term monitoring related to climate change impacts on dryland bryophytes including biocrust species.

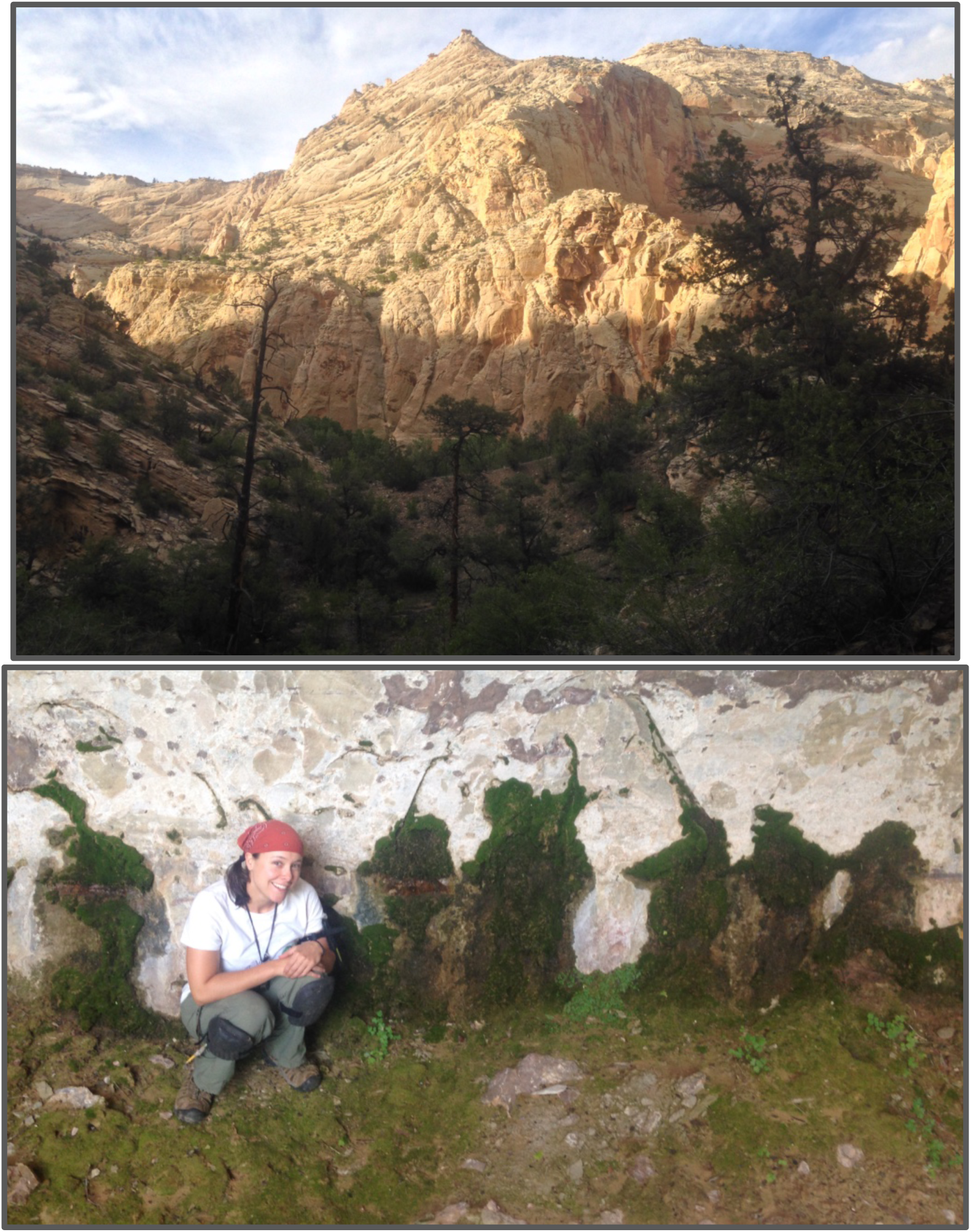

**Cover photos** (by T. A. Clark): View of sandstone canyon wall along the Escalante River Trail taken during a July collection trip in 2015 (top) during which riparian bryophytes were collected by authro, T. A. Clark, (shown in photo) at a sandstone seep (bottom).

**Bureau of Land Management’s National Landscape Conservation System Grant Cooperative Agreement #L14AC00275 issued to P.I. Lloyd R. Stark, UNLV**

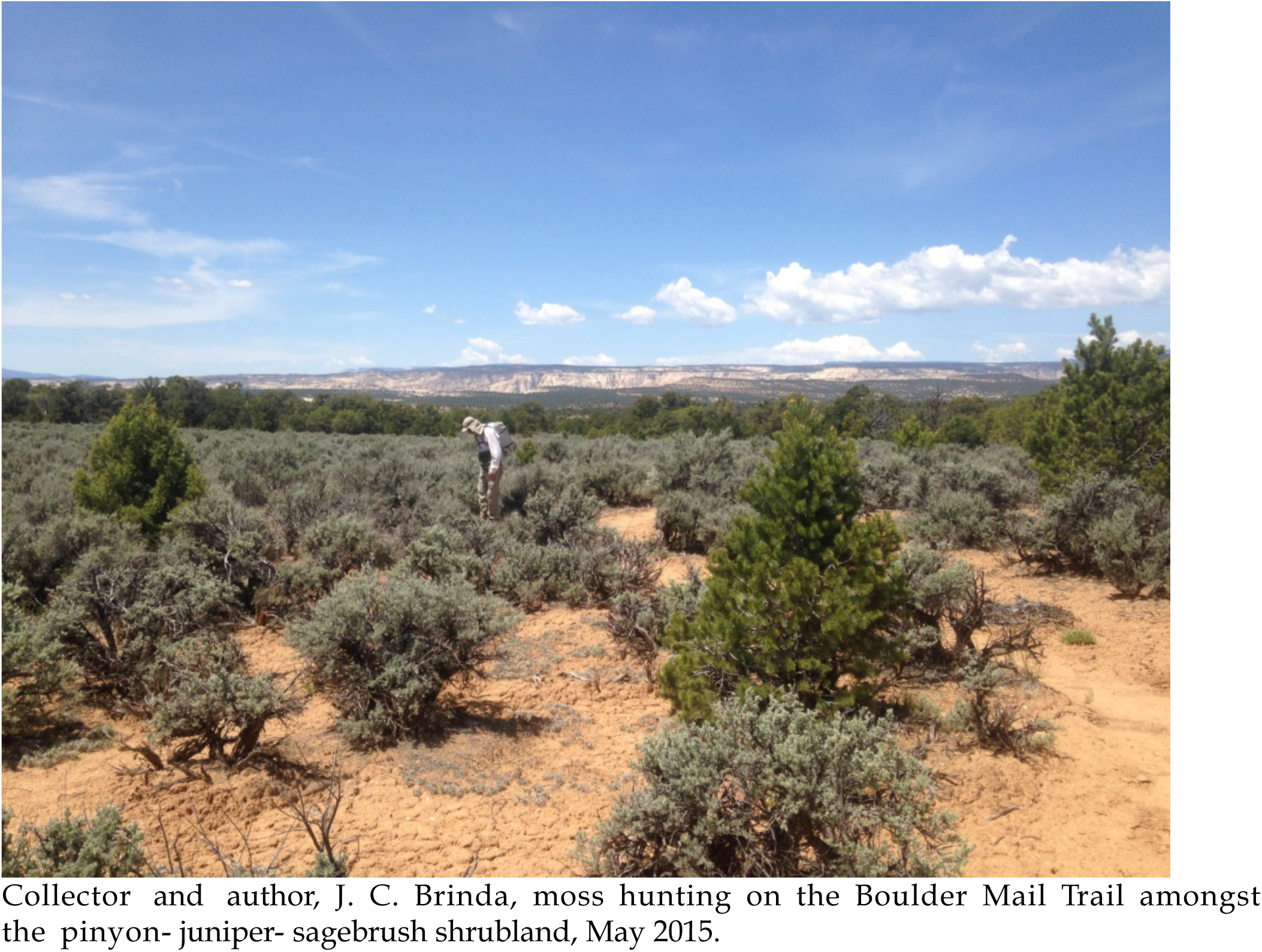

**Copyright 2020 by the authors:** All content contained herein is the property of the authors and all images the property of Theresa A. Clark and should not be used without permission except for education, in which case inclusion of the author/photographer’s name in citation or superimposed over any image(s) is requested.

## Introduction

Although bryophytes (mosses, liverworts, and hornworts) are generally not common or abundant in arid and semi-arid regions, past collecting in the American Intermountain West and Southwest continues to document state and national species records including while also uncovering species new to science (Spence 1987, 1991, Stark et al. 2002, Spence 2007, Brinda et al. 2007, Clark 2012, Brinda et al. 2014). Recent surveys of Grand Canyon-Parashant National Monument (GCPNM, Stark and Brinda 2011) and Grand Canyon National Park (GCNP, Clark 2012) have demonstrated that bryophyte floras (i.e. bryofloras) on the Colorado Plateau can be extremely diverse. Together, these inventories revealed over 50 new species records for Arizona and Nevada and at least four species new to science (authors Brinda, Clark, & Spence, *unpublished data*).

As the largest protected public land on the Colorado Plateau, GSENM encompasses a broad range of geology, soils, topography, and associated micro- and meso-climates that should also foster a large bryoflora comparable to that of GCNP and GCPNM. Protection of the diverse natural resources in GSENM was one of the primary reasons for the monument’s designation in 1996 by President Bill Clinton. Although Nevada has a recently published list of bryophytes (Brinda et al. 2007), Utah still lacks a modern list, with the last formal checklist produced over 30 years ago (Flowers 1961 & 1973, Spence 1988). Moreover, no bryophytes have been collected inside the GSENM, despite the importance this monument has placed on biological soil crust (biocrust) conservation and ecological research (e.g. Bowker et al. 2006). However, a small riparian area adjacent to GSENM, in Deer Creek, revealed two moss species new to Utah, *Anomobryum concinnatum* and *Rhynchostegium aquaticum*, which represented significant and unexpected range extensions for these species (Fertig et al. 2009, 2013).

The BLM-GSENM Proclamation identifies one type of community supporting bryophytes, the biological soil crusts (biocrusts), as “protected objects” recognized for their important ecosystem services. Mosses in biocrust help absorb and retain water, stabilize soils, aid nutrient cycling, fix carbon, and assist seedling germination (Bowker et al. 2006). Large expanses of the Colorado Plateau landscape support extensive populations of two arid-adapted moss species, *Syntrichia caninervis* and *S. ruralis*, which grow on exposed soil or as keystone members of biocrust communities. The ecosystem services of bryophytes extend to other substrates including rock and riparian habitats (Glime 2015). Species in the rock moss genus, *Grimmia*, are common on all rock types throughout drylands in North America, aiding dust capture, nitrogen fixation (indirectly by supporting cyanobacterial species), water retention, and stable-habitat creation for microbes and invertebrates (Glime 2014). Aquatic bryophytes found in streams and associated with springs are also important components of the food base for invertebrates and fish (Glime 2014).

However, global climate change threatens many of these bryophytes and their ecosystem functions. The American Southwest, including portions of the southwestern Colorado Plateau, is experiencing some of the most rapid increases in temperature in the U.S. linked to recent, large-scale die-offs in dominant vegetation; studies have already begun to document changes in the composition of vascular plants as a result of combined drought and higher temperatures (Breshears et al. 2005; Munson et al. 2011). In order to document similar, predicted changes in bryophyte communities, research must prioritize baseline inventories throughout the American Southwest where such climate and vegetation changes are likely to have compounding impacts on bryophytes, many of which already exist at or near their physiological limits.

To their detriment, bryophytes have been shown to serve as excellent indicators of early climate change due to their sensitive water relations and required microclimate conditions, including substrate chemistry, humidity, precipitation (timing and amount), and insolation (Gignac 2001, Tuba et al. 2011). Drastic range alterations have already been documented for several bryophytes as certain high-risk populations have disappeared from prior regions while new areas are colonized as their conditions become suitable with climate change (Tuba et al. 2011). However, recent field and laboratory studies suggest that even widespread, stress-tolerant species such as *Syntrichia caninervis* in the Mojave Desert, appears highly susceptible to subtle changes in temperature, precipitation, UV, and humidity (Barker et al. 2005; Slate et al. 2019, Tuba et al. 2011, Coe et al. 2020). Furthermore, aquatic and semiaquatic bryophytes are particularly vulnerable to drought-induced declines in water availability, with local extinction a likely outcome, in response to the predicted increasing frequency of long-term drought. Changes in the distribution and abundance of such sensitive hydric bryophytes could have a large impact on regional diversity as streams, springs, and seeps are obligate habitat for many Southwestern species (Clark 2012).

With both xeric and hydric bryophytes appearing susceptible to climate change in drylands, there is a clear and urgent need to expand the baseline inventory of Southwestern bryophytes.

## Objectives

The primary objective of this research was to complete a preliminary bryoflora for Grand Staircase-Escalante National Monument (GSENM) targeting topographically-buffered sites predicted to be potential diversity hotspots while collecting at more exposed sites (in-transit to target sites) to capture site-level bryophyte richness and composition at habitats with reduced shade and moisture where lower diversity was predicted. With this targeted approach, the aim was to capture ca. 80% of the local species richness while producing a preliminary circumscription of common species distributions throughout the monument. The second objective was to produce a qualitative ecological assessment of patterns in bryophyte diversity across the monument speculating environmental drivers of diversity hotspots (e.g. microhabitat diversity, elevation, vegetation type, and topography). The third goal was to make a qualitative comparison between gamma diversity (landscape level species richness) and community composition of GSENM and Grand Canyon National Park focusing on compositional trends by bryophyte growth form and taxonomic levels (e.g. families and genera).

## Methods

### Site selection

Because bryophytes occur in low abundance and frequency in drylands, site selection for floristic sampling was exploratory rather than random. We used maps of the monument to target potential diversity hotspots based on the ecological principles of habitat buffering (Shi et al. 2014) choosing sites across the elevation gradient and near perennial streams and tall, north-facing topographical features including canyon walls, plateau cliffs, and hillslopes that were visible on 1:75000-scale topo maps or on-site during explorative travel through the monument.

### Floristic habitat sampling

Collection sites were searched systematically for bryophytes within a ca. 0.25-mile radius except for narrow macrohabitats like gorges in which samples span longer distances along the channel (**Table 1**). Collections were made at over 40 localities across the three ecological study units of the monument (**Fig. 1**): The Staircase, Kaiparowits Plateau, and Escalante Canyons (**Table 1**). The sampling protocol we implemented is called floristic habitat sampling, which has been shown in several bryophyte studies to maximize local species capture when compared to plot-based sampling (e.g. Newmaster et al. 2005). In this approach, all suitable substrates (**Fig. 2**) are searched for bryophytes at each site while strategically targeting common and unique microhabitats. This method enables discovery of both common and rare species at a site, many which would go uncovered with randomized plot sampling at any scale.

**Figure 1.**
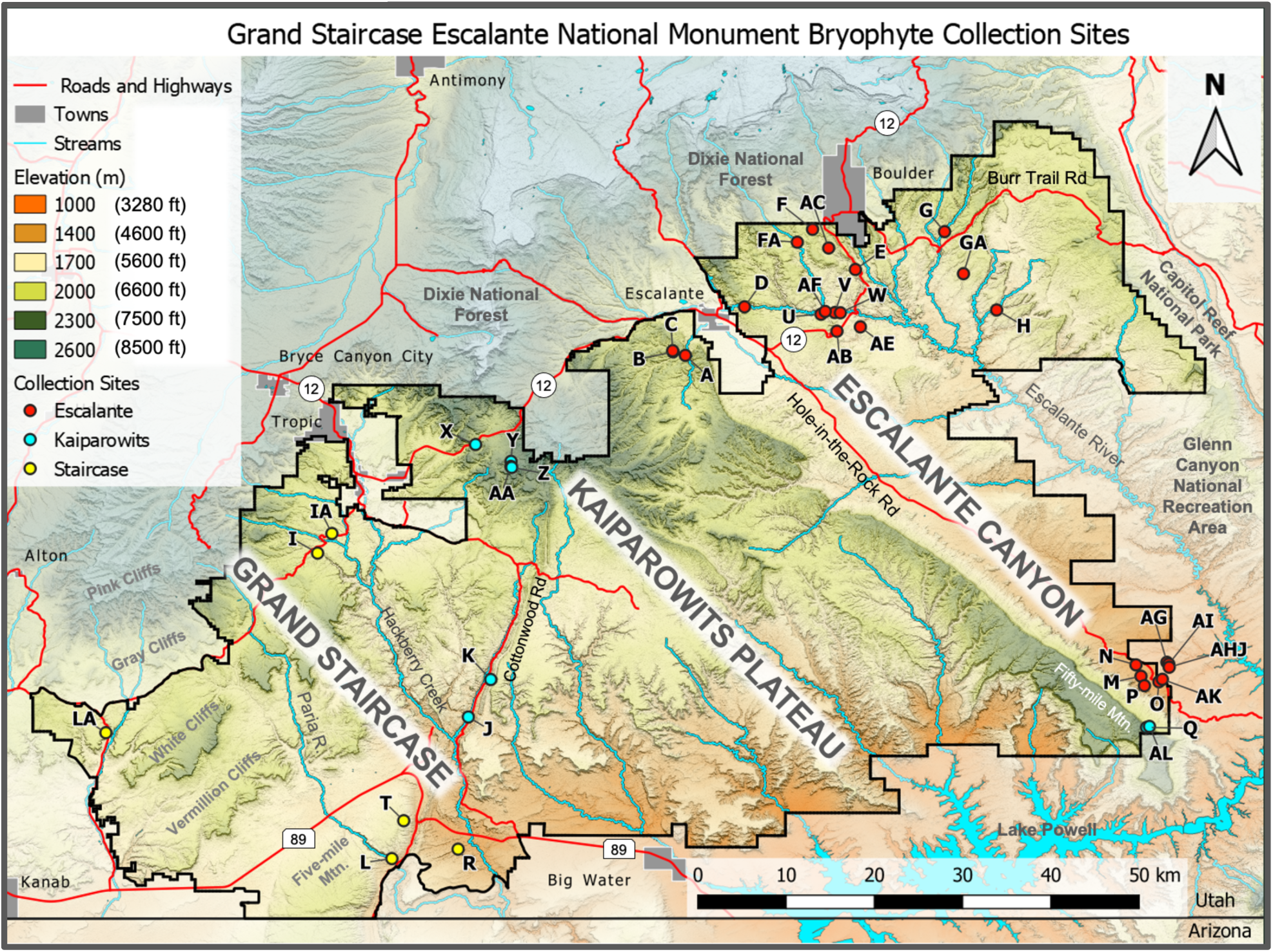
Location of 40 collection sites sampled in 2015 & 2016 within the original boundaries of GSENM, Utah. Elevation bands are color-coded. See also **Table 1**.

**Figure 2.**
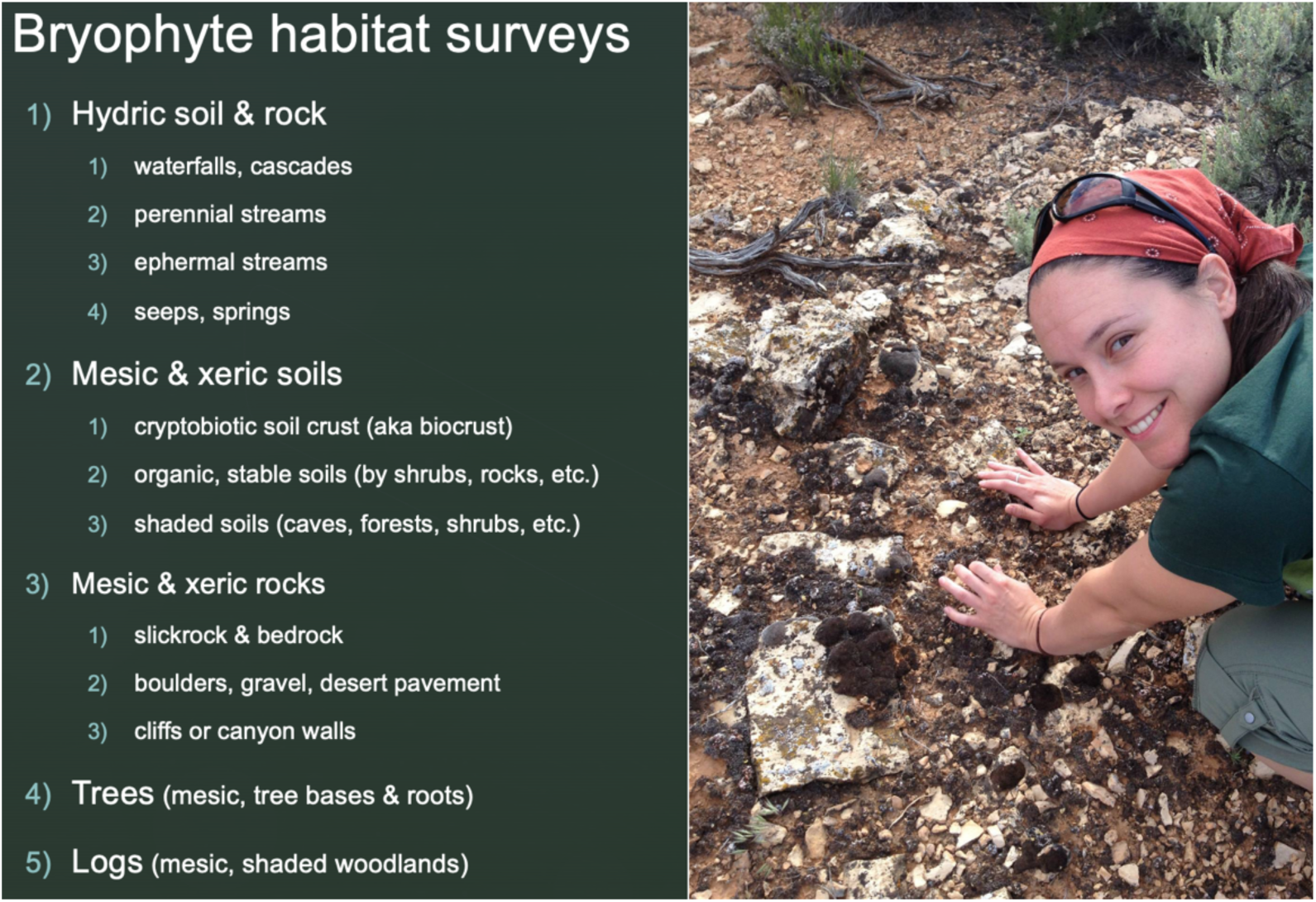
Floristic habitat sampling surveys were performed at sites by author T. A. Clark (shown) and involved searching all habitats (e.g. waterfalls) and their substrates (e.g. wet rock and soil) known to support bryophytes in drylands.

**Table 1.** Sites sampled for bryophytes in the GSENM (see map **Fig. 1**). *Site* and *Locality* code for the GSENM study regions: Escalante (**red**), Kaiparowits Plateau (**yellow**), and The Staircase (**blue**). Sites are ordered by color-coded classes of *Topography* and *Vegetation*. **Abbreviations**: Trailhead (TH) Hardwood (HW), stream flow (ephemeral/perennial), South- or N-facing (S/N). Additional data are available in **Supplement 3** (**Catalog of Sites**).

| Site | Locality (details) | Topography | Vegetation | Elev (ft) | Locality description | Date | Latitude | Longitude | Collections |
| --- | --- | --- | --- | --- | --- | --- | --- | --- | --- |
| B | Mitchell Canyon (N) | canyon-ephemeral | Mixed Conifer | 6200 | N-facing slope inside Mitchell Canyon near its junction with Coal Bed Canyon. | 5/27/15 | 37.723059 | -111.650461 | 60 |
| X | Henriville Creek | canyon-ephemeral | Mixed Conifer | 6600 | Unnamed canyon SE of Rt 12 & NE of Henriville and E of The Blues. | 5/26/15 & 7/15/15 | 37.605104 | -111.901837 | 78 |
| A | Coal Bed Canyon | canyon-ephemeral | Pinyon-Juniper | 6100 | Coal Bed Canyon between Alvey Wash Rd (#300) & Mitchell Canyon. | 5/27/15 | 37.718696 | -111.634757 | 40 |
| C | Mitchell Canyon (S) | canyon-ephemeral | Pinyon-Juniper | 6200 | Along a S-facing slope of Mitchell Canyon near junction w/ Coal Bed C. | 5/27/15 | 37.7245467 | -111.6509033 | 11 |
| K | Cottonwood Road | canyon-ephemeral | Pinyon-Juniper | 5300 | E of Cottonwood Rd (#400), along small NW-facing draw on The Cockscomb. | 6/4/15 | 37.305659 | -111.882494 | 22 |
| R | Long Canyon | canyon-ephemeral | Pinyon-Juniper | 4500 | Shallow canyon off Rt 89 after the Dolphin fin. | 6/22/15 | 37.089506 | -111.924052 | 12 |
| H | Wolverine Creek | canyon-ephemeral | Riparian | 5300 | Along the canyon formed by Wolverine Creek above its confluence with Horse Canyon. | 6/1/15 | 37.776471 | -111.238527 | 66 |
| L | Buckskin Gulch | canyon-ephemeral | Scrubland | 5000 | Along Buckskin Gulch above House Rock Valley Road (#700). | 6/5/15 | 37.0775 | -112.008346 | 51 |
| AG | Willow Gulch (garden) | canyon-ephemeral | Seep/ spring | 4100 | Hanging garden locality <0.5 mi from Willow Gulch trailhead. | 6/7/16 | 37.3276358 | -111.0218712 | 11 |
| AHJ | Willow Gulch (seep) | canyon-ephemeral | Seep/ spring | 4100 | Seeping canyon bottom, locality <2 mi from road TH. | 6/7/16 & 6/9/16 | 37.3213651 | -111.0189831 | 9 |
| AI | Willow Gulch (gorge) | canyon-ephemeral | Seep/ spring | 4100 | <30 min hike from TH to narrow (<30 m wide) portion of gorge. | 6/8/16 | 37.3254361 | -111.019502 | 30 |
| AK | Willow Gulch (SE finger) | canyon-ephemeral | Seep/ spring | 4200 | SE finger of Willow Gulch near the GSENM boundary. | 6/9/16 | 37.3059667 | -111.0276833 | 17 |
| V | Escalante Natural Bridge | canyon-perennial | HW-Pinyon-Juniper | 5300 | Cavernous natural bridge area <0.25 mi off the Escalante River Trail. | 7/13/15 | 37.773235 | -111.444451 | 39 |
| AF | Escalante Canyon | canyon-perennial | Pinyon-Juniper | 5300 | Basalt deposit trailside along the Escalante River Canyon Trail. | 6/6/16 | 37.7746551 | -111.4567478 | 15 |
| AC | Upper Calf Creek Falls | canyon-perennial | Riparian | 6000 | Along the Upper Calf Creek trail with deep pools and waterfalls. | 8/13/15 | 37.855433 | -111.452335 | 13 |
| AE | Phipps Wash | canyon-perennial | Riparian | 5300 | E-W running wash off Escalante R just S of Rt 12 before Maverick Bridge. | 8/15/15 | 37.75494 | -111.411974 | 30 |
| E | Lower Calf Creek Falls | canyon-perennial | Riparian | 5500 | Along small tributary to Calf Creek, southeast of Lower Calf Creek Falls | 5/29/15 | 37.828056 | -111.418366 | 88 |
| U | Escalante Canyon | canyon-perennial | Riparian | 5300 | Escalante Canyon Trail south of the Natural Bridge | 7/12/15 | 37.771027 | -111.462066 | 30 |
| W | Escalante River | canyon-perennial | Riparian | 5200 | Escalante R. riparian, off Rt 12, E of Escalante Natural Bridge near trail. | 7/14/15 | 37.772918 | -111.437393 | 19 |
| D | Escalante Canyon Trail | canyon-perennial | Riparian, HW-Conifer | 5800 | 2-hr hike from TH to a N-facing bend in Upper Escalante Canyon between Escalante village and Mamie Creek | 5/28/15 | 37.780508 | -111.559461 | 82 |
| FA | Sand & Sweetwater Creeks | canyon-perennial | Riparian, HW-Conifer | 6200 | Off Rt 12 in Sugarloaf region; steep trail to confluence of Sand and Sweetwater Creeks. | 5/30/15 | 37.862836 | -111.492268 | 50 |
| G | The Gulch | canyon-perennial | Riparian, HW-Conifer | 5800 | Along The Gulch (steep ravine) between Steep Creek and Water Canyon. | 5/31/15 | 37.876319 | -111.304807 | 62 |
| GA | Water Canyon | canyon-perennial | Riparian, HW-Conifer | 6000 | Along Water Canyon near its confluence with The Gulch. | 5/31/15 | 37.8227048 | -111.2807861 | 30 |
| I | Bull Gorge | canyon-perennial | Riparian, HW-Conifer | 6200 | Along Bull Valley Gorge below Skutumpah Road (#500). | 6/3/15 | 37.466711 | -112.102951 | 87 |
| J | Lower Hackberry Canyon | canyon-perennial | Riparian, HW-Conifer | 4800 | Along Lower Hackberry Canyon with shallow stream and Cottonwoods. | 6/4/15 | 37.257905 | -111.911012 | 45 |
| IA | Bull Gorge Upland | flat | Pinyon-Juniper | 6000 | Near the entrance to Bull Valley Gorge above Skutumpah Road (#500). | 6/3/15 | 37.491683 | -112.084751 | 11 |
| LA | Johnson Canyon Rd. | flat | Scrubland | 5900 | At base of N-facing wall off Rd. #603 in Johnson Canyon just east of Mark Point. | 6/16/15 | 37.237909 | -112.372495 | 4 |
| M | Cottonwood Spring | flat | Seep/spring | 4400 | Off Rd #275 in shallow E-W running draw close to Fifty Mile Mountain. | 6/17/15 | 37.309859 | -111.054855 | 5 |
| O | Cave Spring | flat | Seep/spring | 4200 | Cave Springs vicinity below Cave Point near end of Hole-in-the-Rock Rd. | 6/18/15 | 37.30279 | -111.031992 | 11 |
| Y | The Blues (peak) | mountain | HW-Pinyon-Juniper | 7500 | Small mountain top off Mudsprings Rd. | 8/10/15 | 37.583535 | -111.856722 | 11 |
| AA | The Blues (ravine) | mountain | Mixed Conifer | 7500 | Small mountain ravine off Mudsprings Rd. | 8/11/15 | 37.577227 | -111.856913 | 15 |
| Z | The Blues (ravine) | mountain | Mixed Conifer | 7600 | N-facing ravine 1 mi from Mudsprings Rd over steep, mountain terrain. | 8/11/15 | 37.575955 | -111.85584 | 8 |
| AL | Fifty Mile Cliff | plateau | HW-Pinyon-Juniper | 6500 | Along steep trail with huge boulders up the plateau starting at Fifty Mile Bench Rd. trailhead. | 6/10/16 | 37.2459258 | -111.0440942 | 19 |
| Q | Fifty Mile Mountain | plateau | HW-Pinyon-Juniper | 7100 | Atop plateau in a shallow (<200' deep) draw. | 6/19/15 | 37.244666 | -111.047779 | 32 |
| P | Sooner Road | plateau | Pinyon-Juniper | 4600 | Boulder field flat off Sooner Rd. at base of Sooner Slide Rd. | 6/18/15 | 37.297786 | -111.050456 | 4 |
| T | Five Mile Mtn. | plateau | Pinyon-Juniper | 5800 | NW rim of the plateau. | 6/23/15 | 37.125752 | -111.993295 | 14 |
| AB | Escalante Slickrock | slickrock | HW-Pinyon-Juniper | 5600 | Open slickrock region (no trail) immediately off Rt 12, just S of Kiva Coffee. | 8/12/15 | 37.749431 | -111.441807 | 22 |
| F | Boulder Mail Trail | slickrock | Mixed Conifer | 6700 | Small bluff along the Boulder Mail Trail about 0.5 km southwest of McGath Bench Road (#101). | 5/30/15 | 37.879202 | -111.472822 | 37 |
| N | Sooner Rocks | slickrock | Scrubland | 4400 | Off Hole-in-the-Rock Rd. near Dance Hall Rock. | 6/18/15 | 37.324314 | -111.061236 | 26 |

### Species determinations

Taxonomy and state record status follows the Bryophyte Flora of North America almost exclusively (BFNA editors), however we propose alternative treatments when necessary. Over 400 collections made by J. C. Brinda in 2015 have been identified and are referenced with collection numbers [#7564 – 7932] in **Supplement 1** (**Catalog of Specimens**). Determinations for 800 collections made by T.A. Clark in 2015-2016 are in-progress [#1260 – 2105] with tentative vouchers available in **Supplement 2** (**Vouchers Catalog**) to ultimately be deposited in the University of Nevada Las Vegas (***UNLV***) and GSENM-BLM herbaria.

### Specimen metadata & public outreach

Half of the bryophyte collections (480) are currently georeferenced in the iNaturalist Outreach Project (https://www.inaturalist.org/projects/bryophytes-of-grand-staircase-escalante-national-monument) created by T. A. Clark. Locality and habitat data for each collection vary in detail, but all observations include at least the latitude and longitude (via iPhone), date of collection, and a photograph of the specimen in its natural habitat before collection. A subset of the iNaturalist observations (i.e. formal bryophyte collections) at each site were more detailed and included a range of habitat data (e.g. substrate, aspect, slope, vascular plant vegetation, shade level, moisture status, disturbance) along with a photo series capturing macrohabitat, microhabitat, and closeups of the specimen. These data are available to the public as georeferenced observations in the iNaturalist project.

## Results: Preliminary Bryophyte Checklist

This preliminary bryophyte inventory for Grand Staircase Escalante National Monument (GSENM) included over 1000 collections made across 40 localities (i.e. macrohabitat types) spanning local gradients in bryophyte shade and moisture availability (**Table 1**). At present, the growing checklist contains 117 taxa of liverworts and mosses including 27 families, 65 genera, 116 species, 9 varieties, and 1 subspecies. Noteworthy records include 49 putative taxa new for the state of Utah, and 2 undescribed species in the genera *Grimmia* and *Schistidium*. We propose 4 of these species be considered for addition to the recently revised bryoflora of North America.

### Preliminary Annotated List of Taxa

The following alphabetical list of liverworts and mosses in GSENM is preliminary with some collections still under study by the authors and additional specialists. Consequently, the frequency of collections for each taxon will change prior to publication and many collections may reveal additional species records for the park. It is unlikely that taxa listed here will be removed from the final list, although taxonomic revisions continue to progress in bryology and necessitate cross-checking synonyms many of which are included in **Supplement 4**. Selected collection numbers for each author and Dr. Roger Rosentreter are included in brackets and their metadata are available in report Excel files. Voucher specimens (at least two per taxon) from T. Clark are in preparation (**Supplement 2**) and will be delivered to the Kanab BLM Visitor Center. For each taxon, relevant taxonomic, morphological, and distributional data have been summarized in **Table 1** and **Table 2**. Taxa new to Utah (*) were determined from comparison with state distributions in the Bryophyte Flora of North America (BFNA) and include taxa not treated in BFNA, but for which we suggest future consideration for inclusion in the North American bryoflora (**).

**Table 2.** Hotspot ecology, community demographics, and species for the 8 sites having ≥28 species. The number of acrocarpous (***A***), pleurocarpous ***(P***) mosses and liverworts (***L***) are listed. Colors and abbreviations follow that in Table 1.

| Site | Locality (details) | Topography | Vegetation | Elev (ft) | A | P | L | Total spp. | Species (6-letter abbreviations; Sp. = species) |
| --- | --- | --- | --- | --- | --- | --- | --- | --- | --- |
| D | Escalante Canyon Trail | canyon-perennial | Riparian, HW-Conifer | 5800 | 25 | 7 | 2 | 34 | Anohan, Brafen, Bryrec, Cepdiv, Cerpur, Didrig, Ditle, Eucver, Eurpul, Fisbry, Fisdub, Gemcae, Gricar, Gricri, Gripla, Gripul, Hommex, Hyprev, Hypvau, Mniari, Plajun, Pohcru, Psetec, Rebhem, Roslae, Saegla, Schamb, SchSp., Syncan, Synsub, Toralp, Tortor, Torbre, Torobt |
| E | Lower Calf Creek Falls | canyon-perennial | Riparian | 5500 | 17 | 9 | 6 | 32 | Anepin, Anohan, Concom, Consal, Crafil, Didnev, Didtop, Discap, Eucver, Fostex, Funhyg, Griano, Gripul, Hygnot, Hyghum, Hyprev, Hymrec, Imbmue, Jafrau, Leppyr, Leprip, Marpal, Moebly, Pohwah, Psetec, Prequa, Ptymee, Ptypse, Rhyaqu, RosSp., SchSp., Synsub |
| H | Wolverine Creek | canyon-ephemeral | Riparian | 5300 | 30 | 2 | 0 | 32 | Brafen, Bryrec, Didans, Didaus, Didnev, Didrig, Encint, Eucver, Fisbry., Griano, Gricri, Gripla, Griova, Gripul, Grivag, Gymcal, Jafrau, Jafwri, Psetec, Psecr, PteSp., Rostor, Schpul Syncan, Synmon, Toralp, Torbre, Torine, Tormuc, Torobt, Triplan, Weiand |
| FA | Sand & Sweet-water Creeks | canyon-perennial | Riparian, HW-Conifer | 6200 | 23 | 7 | 1 | 31 | Brafen, Brafri, Cepdiv, Concom, Coscal, Didrig, Didtop, Didvin, Ditle, Encint, Eurpul, FisSp., Gemcae, Griano, Gricar, Gricri, Henhei, Hyprev, Jafrau, Leprip, Lopexc, Ortano, PohSp., Psetec, Ptycre, Rosfla, SchSp., Syncan, Synsub, Toralp, Torbre |
| J | Lower Hackberry Canyon | canyon-perennial | Riparian, HW-Conifer | 4800 | 28 | 2 | 0 | 30 | Barriv, CroSp., Didbra, Didrig, Didtop, Didvin, Enchrha, Eucver, Gemcae, Griano, Gricap, Gricri, Griorb, Gripla, Gripul, Gymcal, Henhei, Imbmue, Jafrau, Psetec, Pteova, Ptymee, SchSp., Syncan, Synmon, Synsub, Toralp, Torbre, Tormuc, Torine |
| X | Henrville Creek | canyon-ephemeral | Mixed Conifer | 6600 | 27 | 3 | 0 | 30 | Barbol, Brafen, Bryarg, BrySp., Concom, Croab, Didrig, Encint, Enchrha, Funhyg, Griano, Gemcae, Griano, Gricap, Gripla, HymSp., Hyprev, Orflo, Psetec, Pteova, PtySp., RosSp., SchSp., Syncan, Synmon, Synrur, Synsub, Toralp, Tormuc, Torobt |
| G | The Gulch | canyon-perennial | Riparian, HW-Conifer | 5800 | 23 | 4 | 1 | 28 | Anepin, Barbol, Brafen, Bryarg, Bryrec, Concom, Crafil, Didrig, Didtop, Encint, Eucver, Fisbry, Gemcae, Griano, Gricri, Gripla, Gripul, Henhei, Imbmue, Jafrau, Psetec, Ptymee, Syncan, Synmon, Synsub, Toralp, Tormuc, Torobt |
| V | Escalante Natural Bridge | canyon-perennial | HW-Pinyon-Juniper | 5300 | 17 | 8 | 3 | 28 | Brafen, Braalb, Bryrec, BrySp., Bryarg, Cepdiv, Concom, DidSp., Didvin, Eucver, Eurpul, FisSp., Gemcae, Gemval, GriSp., Hyprev, Marpal, Psetec, PohSp., Potsp, PseSp., HomSp., PtySp., RosSp., SchSp., Syncan, Toralp, Lopexc |

### Liverworts: Phylum Hepaticophyta

(putatively new to *Utah or **new to N. America)

**Aneura pinguis** (L.) Dumort. [Brinda 7729, Clark 1378, 1489]

**\*Cephaloziella divaricata** (Sm.) Schiffn. [Brinda 7668, 7749a, Clark 1348, 1809, 1751]

**Conocephalum salebrosum** Szweyk., Buczk. & Odrzyk. [Brinda 7718, Clark 1380, 1784]

**\*Fossombronia texana** Lindb. [Brinda 7725, 7727, Clark 1400]

**\*Lophoziopsis excisa** (Dicks.) Konstant. & Vilnet [Brinda 7749b, Clark 1807]

**\*Marchantia paleacea** Bertol. [Brinda 7695, 7704, Clark 1394]

**Marchantia polymorpha** subsp. **ruderalis** Bischl. & Boissel.-Dub. [Brinda 7696, Clark 1774]

**\*Moerckia blyttii** (Mørch ex Hornem.) Brockm. [Brinda 7693]

**Preissia quadrata** (Scop.) Nees [Brinda 7697]

**Reboulia hemisphaerica** (L.) Raddi [Brinda 7657]

**Riccia glauca** L. [Clark 1688]

### Mosses: Phylum Bryophyta

**\*Aloini rigida** (De Notaris) Delgadillo-Moya [Clark 1596]

**\*Anoectangium handelii** Schiffn. [Brinda 7659, 7714, 7733; Clark 1392, 1440, 1356]

**Amblystegium serpens** (Hedwig) Bruch [Clark 1797, 1853]

**Barbula bolleana** (Müller Hal.) Brotherus [Clark 1276, 1476]

**Brachytheciastrum collinum** (Schleicher ex. Mull. Hal.) Ignatov & Huttunen [Clark 1727, 1737]

**Brachytheciastrum fendleri** (Sull.) Ochyra & Żarnowiec [Brinda 7877, 7928, 7581, 7582, 7626, 7627, 7678, 7746, 7800, 7823, 7828]

**\*Brachythecium albicans** (Hedwig) Schimper [Clark 1823]

**Brachythecium frigidum** (Müll. Hal.) Besch. [Brinda 7759]

**Brachythecium rivulare** Schimper [Clark 2105]

**Bryoerythrophyllum recurvirostrum** (Hedw.) P.C. Chen [Brinda 7842, 7875, 7620, 7624, 7664, 7798, 7829; Clark 1363, 1421, 1538, 1540, 1558, 1580, 1682, 1802, 1805]

**Bryum argenteum** Hedwig [Clark 1298, 1319, 1425, 1822, 1865, 1897, 1943, 1957, 1971, 1998, 2053, 1309]

**Ceratodon purpureus** (Hedw.) Brid. [Brinda 7603, 7680, Clark 1367, 1427, 1441, 1453]

**Conardia compacta** (Müll. Hal.) H. Rob. [Brinda 7583, 7687, 7689, 7690, 7691, 7758, 7766, 7768, 7778, 7780, 7786; Clark 1270, 1488, 1743, 1835]

**Coscinodon calyptratus** (Drumm.) C. E. O. Jensen [Brinda 7608, 7673, 7736, Clark 1452, 1343, 1429]

**Cratoneuron filicinum** (Hedw.) Spruce [Brinda 7684, 7779, 7788, Clark 1385, 1405]

**Crossidium aberrans** Holz. & E.B. Bartram [Brinda 7908, 7579, Clark 1275]

**\*Crossidium seriatum** H.A. Crum & Steere [Rosentreter 18952]

**\*Crossidium squamiferum** (Viviani) Juratzka [Clark 2040]

**\*Dicranella varia** (Hedw.) Schimp. [Brinda 7883, Clark 1588, 1914]

**Dicranoweisia crispula** (Hedwig) Milde [Brinda 7587, Clark 1900] This specimen may also be called **Hymenoloma mulahaceni** (Höhn.) Ochyra, not currently included in the BFNA.

**\*Didymodon anserinocapitatus** (X.J. Li) R.H. Zander [Brinda 7818c]

**\*Didymodon australasiae** (Hook. & Grev.) R.H. Zander [Brinda 7932b, Clark 1645, 1523, 1698]

**Didymodon brachyphyllus** (Sull.) R.H. Zander [Brinda 7881, 7734, Clark 2047, 1420, 1623, 1779]

**\*Didymodon insulanus** (De Not.) M.O. Hill [Brinda 7863] Treated under *D. vinealis* in BFNA, but we suggest it be treated as a distinct taxon.

**\*Didymodon nevadensis** R.H. Zander [Brinda 7925, 7721, 7817, 7818a, 7819; Rosentreter 18949, Clark 1399, 1655]

**Didymodon rigidulus** var. **gracilis** (Hook. & Grev.) R.H. Zander [Brinda 7616, 7655, 7744, 7803, 7818d, Clark 1302]

**\*Didymodon rigidulus** var. **icmadophilus** (Schimp. ex Müll. Hal.) R.H. Zander [Brinda 7838, 7897, 7572, 7614, 7617, 7628, Clark 2030]

**Didymodon tophaceus** (Brid.) Lisa [Brinda 7900, 7688, 7700, 7711, 7726, 7755, 7764, 7765, 7769, 7777, 7781, Clark 1412, 1435, 1472, 1620]

**Didymodon vinealis** (Brid.) R.H. Zander [Brinda 7873b, 7891, 7924, 7929b, 7722, Clark, 1300, 1361, 1432, 1433]

**Distichium capillaceum** (Hedw.) Bruch & Schimp. [Brinda 7703, Clark 1391, 1781]

**\*Ditrichum flexicaule** (Schwägr.) Hampe [Brinda 7646, 7679, 7743, Clark 1358, 1437]

**Drepanocladus aduncus** [Clark 1850, 1850, 1952]

**\*Encalypta rhaptocarpa** Schwägrichen [Brinda 7865, 7867, 7868, 7927, 7573, 7615a, 7636, 7637b, 7741, 7801, 7802, 7816, Clark 1273, 1451] These plants fit the description of *E. intermedia* Juratzka, treated as a synonym in FNA, but reported under this name by Horton and others. If accepted, this would be a new taxon for North America.

**\*Encalypta rhaptocarpa** var. **trachymitria** (Ripart) Wijk & Margad. [Brinda 7615b, 7637a, Clark 1301, 1625] This variety is not treated in FNA, which has a broad concept of *E. rhaptocarpa*. If accepted, this would be a taxon record for North America.

**Eucladium verticillatum** (Dicks. ex With.) Bruch & Schimp. [Brinda 7860, 7894, 7623, 7639, 7661, 7686, 7716, 7717, 7720, 7767, 7820, 7832, Clark 1520, 1622, 1681, 1775]

**Eurhynchiastrum pulchellum** (Hedw.) Ignatov & Huttunen [Brinda 7640, 7649, 7652, 7660, 7676, Clark 1346, 1436, 1803, 1803, 1811]

**Fissidens bryoides** Hedwig [Brinda 7862, 7621, 7647, 7775, 7799, 7814, 7833, Clark 1333, 1364] The variety **pusillus** (Wilson) Pursell is a synonym in FNA, but accurately describes the plants.

**\*Fissidens dubius** P. Beauv. [Brinda 7643]

**Funaria hygrometrica** Hedw. [Brinda 7705, Clark 1272, 1413, 1683, 1691]

**\*Gemmabryum badium** (Brid.) J.R. Spence [Brinda 7630]

**Gemmabryum caespiticium** (Hedw.) J.R. Spence [Brinda 7911, Clark 1447, 1485, 1494, 1595, 1602]

**Gemmabryum kunzei** (Hornsch.) J.R. Spence [Clark 1339]

**Grimmia alpestris** (Weber & Mohr) Schleicher [Clark 1721]

**Grimmia anodon** Bruch & Schimp. [Brinda 7851, 7870, 7901, 7903, 7904a, 7567, 7591, 7611, 7737, 7740, 7807, 7812, Clark 1261, 1268, 1288, 1314]

**\*Grimmia capillata** De Not. [Brinda 7888, 7568, 7590, Clark 1287] Not treated in FNA; the plants are either this species or undescribed and should be a new taxon record for North America.

**Grimmia crinitoleucophaea** Cardot [Brinda 7879, 7893, 7902, 7914, 7594, 7653, 7724, 7747, 7751, 7772, 7773a, 7793a, 7808, Clark 1498, 1591, 1619, 1936]

**Grimmia montana** var. **brachyodon** (Austin) Lesq. & James [Brinda 7923, Clark 1651]

**Grimmia orbicularis** Bruch ex Wilson [Brinda 7885, 7915, 7916, 7918, Clark 1598, 1637, 1760, 2064]

**Grimmia ovalis** (Hedw.) Lindb. [Brinda 7809, Clark 1315, 1958]

**Grimmia plagiopodia** Hedw. [Brinda 7849, 7850, 7852, 7854, 7859, 7904b, 7569, 7592, 7593, 7604, 7607, 7644, 7674, 7731, 7774, 7793b, Clark 1368, 1516, 1532, 1553]

**Grimmia pulvinata** (Hedw.) Sm. [Brinda 7846, 7886, 7890, 7892, 7913, 7922, 7605, 7672, 7761, 7810, Clark 2026, 1409, 1529, 1613]

**\*Grimmia vaginulata** Kellman [Brinda 7919, 7921, 7610, Clark 1550, 1669, 1693, 1708]

**\*\*Grimmia sp. nov.** Brinda [Brinda 7857, 7796] Odd falcate-leaved plants belonging to the subgenus Gasterogrimmia (BFNA), but appear undescribed.

**\*Gymnostomum calcareum** Nees & Hornsch. [Brinda 7821, Clark 1522, 1618, 2017, 2049]

**Hennediella heimii** (Hedw.) R.H. Zander [Brinda 7898, 7760, 7785, Clark 1610, 1665, 1680, 1699]

**\*Homomallium mexicanum** Cardot [Brinda 7656, Clark 1814]

**\*Hygroamblystegium varium** subspecies **noterophilum** Sullivant & Lesquereux) Vanderp. & Hedenas [Brinda 7710, 7715, Clark 1389, 1750, 1953, 1783]

**Hygroamblystegium varium** var. **humile** (P. Beauv.) Vanderp. & Hedenäs [Brinda 7706, Clark 1953]

**Hymenostylium recurvirostrum** (Hedw.) Dixon [Brinda 7685, 7719, Clark 1847, 1856, 2009, 2062]

**Hypnum revolutum** (Mitten) Lindberg [Brinda 7834, 7843, 7574, 7613, 7632, 7645, 7671, 7757, 7804, Clark 1359, 1381, 1454, 1514, 1564]

**Hypnum vaucheri** Lesquereux [Brinda 7841, 7869, 7926, 7930, 7934, 7677, Clark 1336, 1636]

**\*Imbribryum muehlenbeckii** (Bruch & Schimp.) N. Pedersen [Brinda 7692, 7708, 7709, 7712, Clark 1408, 1460, 1471, 1609]

**Jaffueliobryum raui** (Austin) Thér. [Brinda 7836, 7920, 7595, 7596, 7597, 7598, 7606b, 7723, 7730, 7754, 7762, 7773b, Clark 1296, 1311, 1411, 1650]

**\*Jaffueliobryum wrightii** (Sull.) Thér. [Brinda 7880, Clark 1293, 1534]

**Leptobryum pyriforme** (Hedw.) Wilson [Brinda 7728, Clark 1373, 1859]

**Leptodictym riparium** (Hedwig) Warnstorf [Clark 1402, 1434, 1793, 1855]

**Leskeaceae** sp. Schimper [Clark 1827]

**Mnium arizonicum** J.J. Amann [Brinda 7666, Clark 1362] **Orthotrichum anomalum** Hedw. [Brinda 7738, 7739] **Orthotrichum diaphanum** Brid. [Brinda 7589] **Orthotrichum flowersii** Vitt [Brinda 7565]

**\*Platydictya jungermannioides** (Brid.) H.A. Crum [Brinda 7662b]

**\*Pleurochaete luteola** (Bescherelle) Theriot [Clark 1929, 1932]

**Pohlia cruda** (Hedw.) Lindb. [Brinda 7669, 7670, Clark 1345]

**Pohlia wahlenbergii** (F. Weber & D. Mohr) A.L. Andrews [Brinda 7702, Clark 1382]

**Pseudocrossidium crinitum** (Schultz) R.H. Zander [Brinda 7818b, Clark 1539]

**Pseudoleskeella tectorum** (Funck ex Brid.) Kindb. ex Broth. [Brinda 7837, 7840, 7878, 7917, 7585, 7586, 7641, 7682, 7750b, 7795, 7830a, Clark 1330, 1340, 1376, 1417]

**Pterygoneurum ovatum** (Hedw.) Dixon [Brinda 7931, 7578, Clark 1267, 1590, 1603, 1660]

**Pterygoneurum subsessile** (Brid.) Jur. [Brinda 7912, 7735, Clark 1424, 1635, 1761, 1942]

**Ptychostomum lonchocaulon** (Müll. Hal.) J.R. Spence [Brinda 7929a]

**\*Ptychostomum meesioides** (Kindb.) J.R. Spence [Brinda 7899, 7694, 7698, 7699, 7707, 7763, 7770, 7783, 7787]

**Ptychostomum pseudotriquetrum** (Hedw.) J.R. Spence & H.P. Ramsay ex Holyoak & N. Pedersen [Brinda 7701, 7782, 7784]

**Rhynchostegium aquaticum** A. Jaeger [Brinda 7713, Clark 1386] Notably, aquatic specimens of *Rhynchostegium* have been historically named *R. riparioides* (Hedwig) Cardot. a species currently not circumscribed for N. America.

**\*Rosulabryum flaccidum** (Brid.) J.R. Spence [Brinda 7753, Clark 1312, 1442, 1571]

**Rosulabryum laevifilum** (Syed) Ochyra [Brinda 7853, 7855, 7662a, 7663]

**\*Rosulabryum torquescens** (Bruch & Schimp.) J.R. Spence [Brinda 7827, Clark 1312, 1442, 1571]

**\*Saelania glaucescens** (Hedw.) Broth. [Brinda 7667, Clark 1351]

**\*\*Schistidium ambiguum** Sull. [Brinda 7675] This species is not treated in BFNA; however, the type specimen is from New Mexico and may justify its inclusion in N. America.

**\*Schistidium flaccidum** (De Not.) Ochyra [Brinda 7612, 7811b]

**\*\*Schistidium sp. nov.** H.H. Blom [Brinda 7844, 7856, 7858, 7889, 7650, 7752] These specimens represent an undescribed species per communication with Hans H. Blom (e.g. Blom & Darigo 2009).

**Syntrichia caninervis** Mitt. [Brinda 7884, 7907, 7909, 7933, 7601, 7631a, 7745, 7771, 7789, 7806; Clark 1297, 1306, 1322, 1338, 1448, 1492, Rosentreter 18948]

**\*Syntrichia montana** Nees [Brinda 7847, 7848, 7584, 7609, 7638, 7792, 7794, 7831, Clark 1887, 1927, 1980, 1995]

**Syntrichia ruralis** (Hedw.) F. Weber & D. Mohr [Brinda 7906, 7631b, Clark 1269, 1416]

**\*\*Syntrichia subpapillosissima** (Bizot & R.B. Pierrot ex W.A. Kramer) M.T. Gallego & J. Guerra [Brinda 2257, 7874, 7876, 7896, 7564, 7575, 7576, 7602, 7629, 7633, 7635, 7681, 7742, 7756, 7776, Clark 1552, 1601, 1654, 1672, 1894] Not treated in FNA, but recent collections by many bryologists in the Southwest reveal this species is more common than *S. ruralis* in montane habitats (a publication on this topic will be cited at https://3dmoss.berkeley.edu/).

**Tortella alpicola** Dixon [Brinda 7835, 7839, 7861, 7570, 7571, 7577, 7588, 7599, 7622, 7658, 7748, 7791, 7805, 7815, Clark 1615, 1828, 1284, 1352]

**\*Tortella inclinata** var. **densa** (Lorentz & Molendo) Limpr. [Brinda 7732] Determined by BFNA taxonomist, Patricia Eckel.

**Tortella tortuosa** (Schrad. ex Hedw.) Limpr. [Brinda 7864, 7665, Clark 1357, 1567]

**Tortula atrovirens** (Turner ex Sm.) Lindb. [Brinda 7618]

**Tortula brevipes** (Lesq.) Broth. [Brinda 7871, 7872, 7873a, 7932a]

**\*\*Tortula brevissima** Schiffn. [Brinda 7887, 7895, 7683, 7750a, 7830b] Not treated in FNA, this is probably an undescribed species but is being reported under this name in North America.

**\*Tortula guepinii** Bruch & Schimp. [Brinda 7929c]

**Tortula inermis** (Brid.) Mont. [Brinda 7910, 7822, Clark 1531, 1626]

**Tortula mucronifolia** Schwägr. [Brinda 7866, 7905, 7580, 7619, 7634, 7797, 7824, 7826, Clark 1290, 1517, 1537, 1556]

**Tortula obtusifolia** (Schwägr.) Mathieu [Brinda 7845, 7566, 7606a, 7642, 7648, 7651, 7654, 7790, 7811a, Clark 1415, 1575]

**\*Trichostomum planifolium** (Dixon) R. H. Zander [Brinda 7882, 7825]

**Pottiopsis sweetii** (E.B. Bartram) Ros & O. Werner has been proposed as a new classification for this species. No collections of *T. planifolium* have been verified in BFNA from Utah, however the synonymized type specimen of *Weissia perligulata* was collected in Utah (BFNA).

**\*\*Weissia andrewsii** E.B. Bartram [Brinda 7600, 7625, 7813, Clark 1294, 1327, 1524, 1557] Treated under *W. controversa* in FNA but is a distinct taxon, compare with the treatment in the moss flora of Mexico.

## Discussion (manager’s summary)

### Bryophyte Composition & Diversity Patterns in GSENM

#### Community composition

##### Growth forms

As expected for arid and semiarid environments, the bryophyte species of GSENM are predominantly mosses (107 species, 91%), which have generally much greater desiccation tolerance than liverworts (11 species, 9%) and hornworts; the latter group may be found in this region of southern Utah, but relatively few hornworts are documented for the American Southwest (Spence et al. 2006, Blisard & Kleinman 2015). Of the mosses, acrocarpous mosses (i.e. those species having vertical shoots with apical sporophytes, **Fig. 3a**) are the most diverse in GSENM (88 species), while the less desiccation-tolerant pleurocarpous mosses (i.e. those species having crawling shoots with lateral sporophytes, **Fig. 3c**) included 19 species (**Supplement 4**).

**Figure 3.**
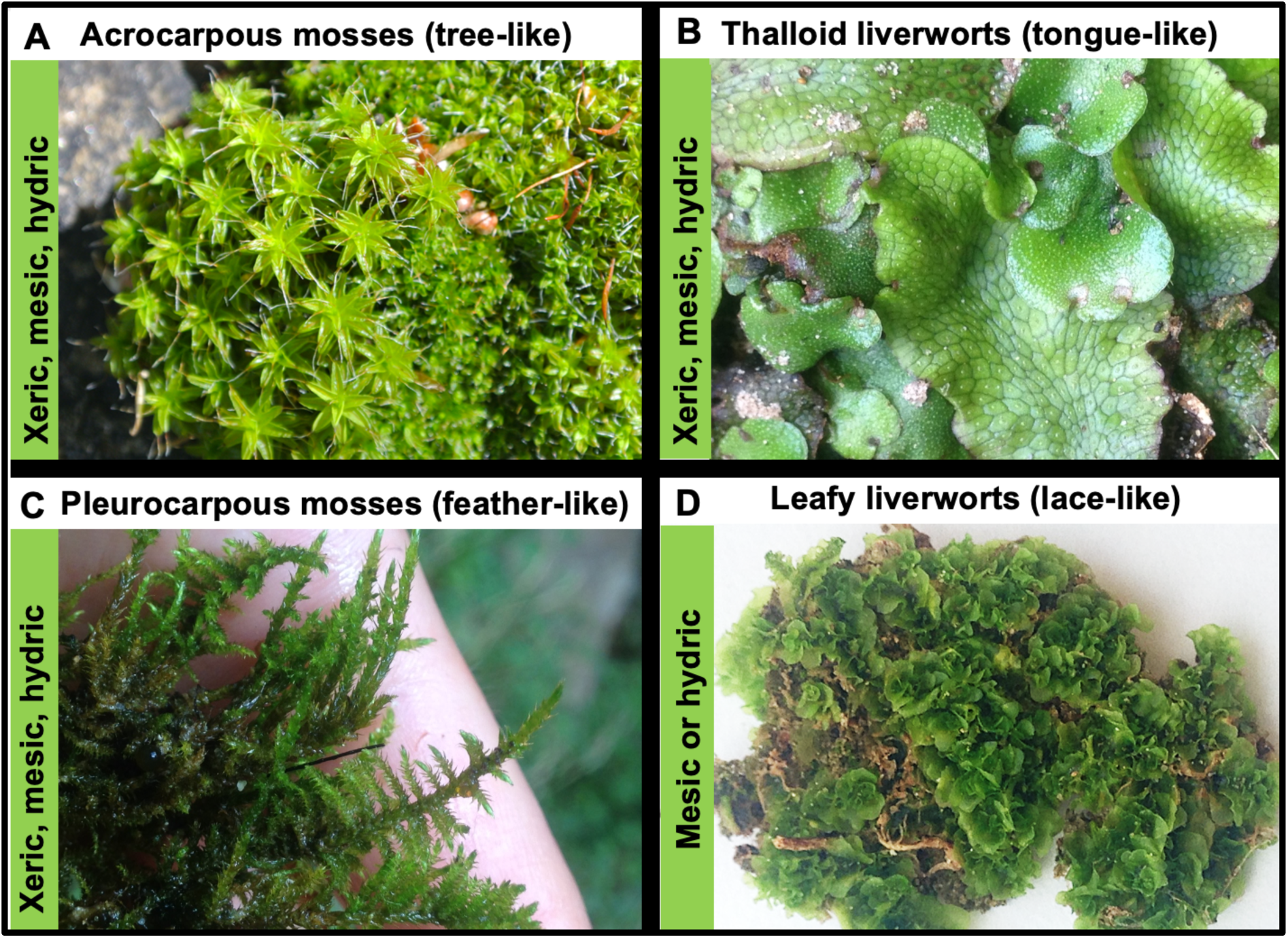
Moisture requirements (xeric, mesic, or hydric) of the four growth forms of bryophytes in GSENM. Charismatic examples of each group in GSENM include acrocarpous mosses in the *Syntrichia* complex (**A**), aquatic pleurocarpous mosses like *Cratoneuron filicinum* (**C**), large riparian thalloid liverworts like *Conocephalum* and *Marchantia* (**B**), and small, delicate leafy liverworts like *Fossombronia* (**D**).

For liverworts, the thalloid growth form was most diverse (8 species, **Fig. 3b**, **Fig. 4**), while the desiccation-sensitive leafy liverworts were rare (3 species, **Fig. 3d**, **Fig. 4**). Leafy liverworts have a crawling, lace-like morphology with stems that adhere closely to their substrates for moisture acquisition and retention. Leafy liverworts are extremely small and rare in drylands like GSENM, while thalloid liverworts are much larger and desiccation tolerant with a thick, flattened strap- or tongue-like morphology – the thallus – which is often forked (**Fig. 3b**). Collectively, the bryoflora of GSENM is comprised of predominantly acrocarpous moss species (75%) followed by pleurocarpous mosses (16%), thalloid liverworts (7%), and leafy liverworts (2%, **Fig. 4**). This species composition strongly matches the composition of bryophytes in Grand Canyon National Park (GCNP), which was 74% acrocarpous and 19% pleurocarpous mosses, and 7% liverworts (based on >1500 collections, Clark 2012). This congruency strongly suggests that additional bryophyte collecting will not change the growth-form composition of the bryoflora in GSENM even as new local species records are added.

**Figure 4.**
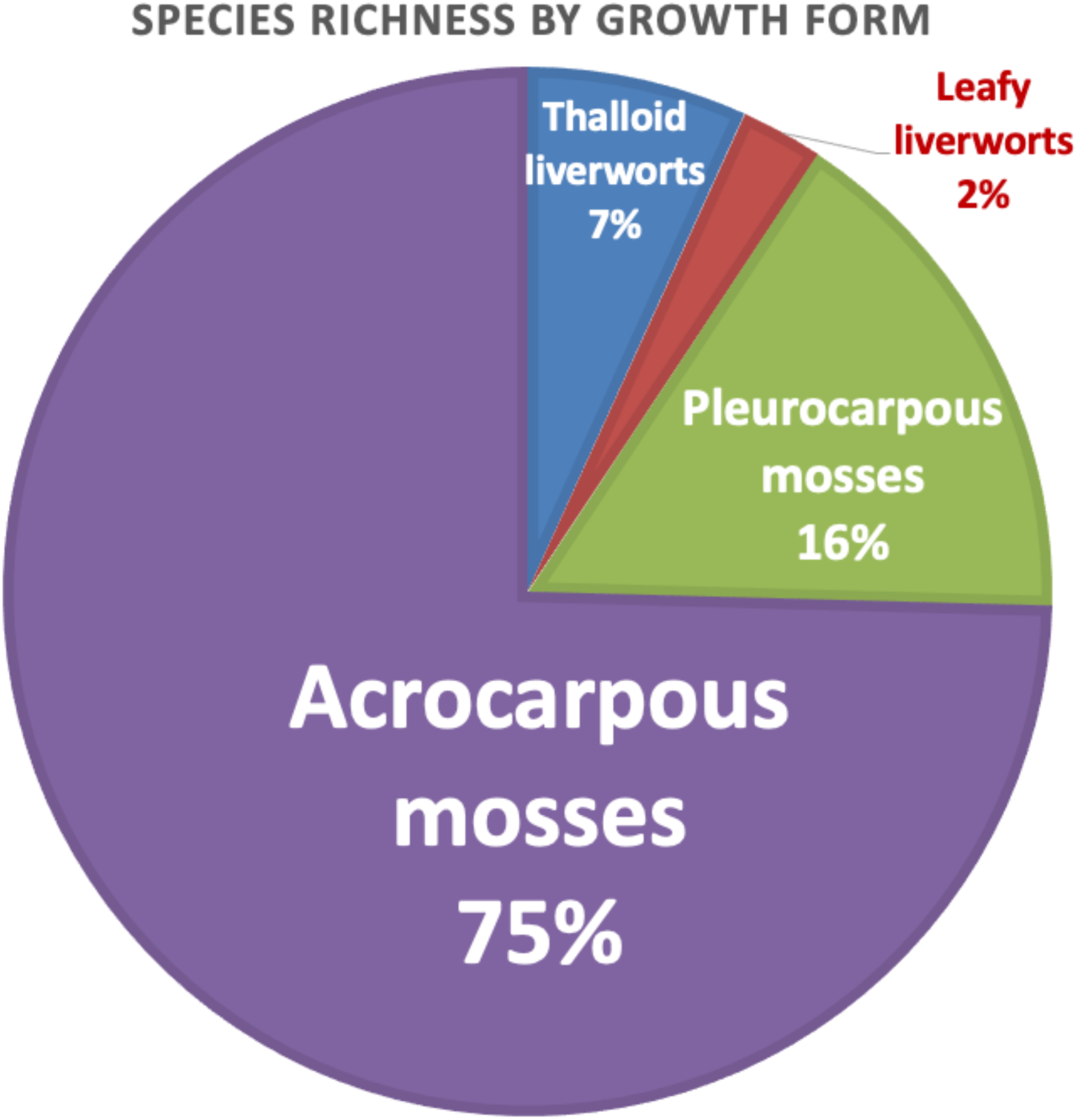
Distribution of 117 bryophyte species found in GSENM by moss and liverwort growth form. The composition closely matches that of Grand Canyon National Park.

##### Bryophyte family diversity

The most diverse families (**Fig. 5**) included highly xeric-soil acrocarpous mosses in the Pottiaceae (35%) and xeric-rock acrocarpous mosses in the Grimmiaceae (15%). Both xeric and mesic species were recovered in the Bryaceae (10% of species) while the pleurocarpous Amblystegiaceae included mesic and hydric species (7%). Therefore, although the acrocarpous mosses of GSENM are the most species-rich group, they may not support a proportional amount of genetic diversity because the 88 acrocarpous species primarily belong to only three families. Nonetheless, there are over two-folds more acrocarpous families (12) than pleurocarpous families (5), strongly suggesting that most of the genetic diversity in the bryoflora is contributed by the acrocarpous mosses (**Supplement 4**).

**Figure 5.**
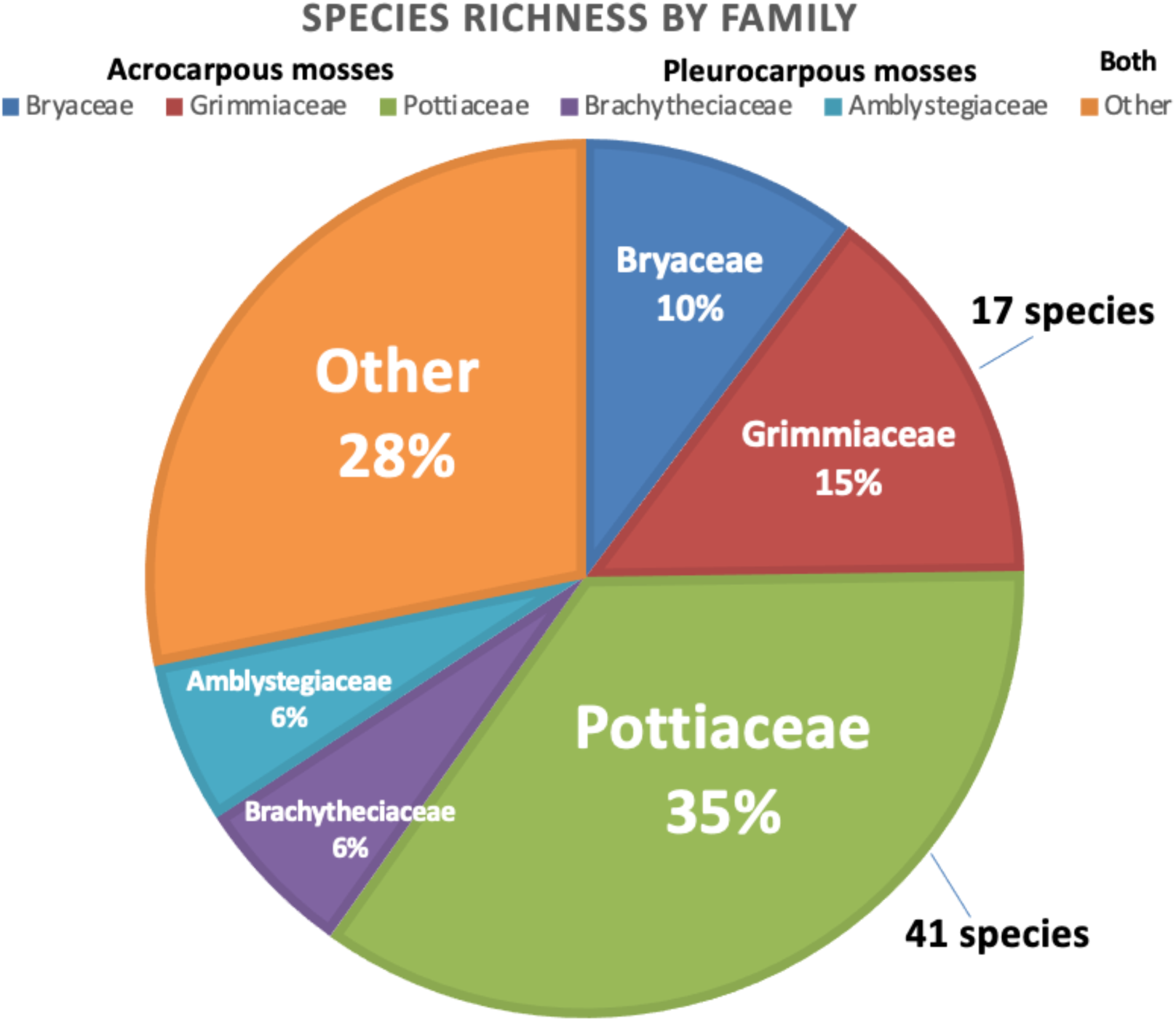
Distribution of 117 GSENM bryophyte species across 27 taxonomic families. Liverworts and mosses comprise the remaining families in *Other*.

Notably, these family demographics closely match that of GCNP, which supports 155 taxa distributed across the Pottiaceae (32%), Grimmiaceae (12%), Bryaceae (11%), Brachytheciaceae (7%), and Amblystegiaceae (3%, Clark 2012). As with growth form demographics, this congruency with GCNP suggests that bryoflora composition may not significantly change as additional species are discovered in GSENM.

### Habitat specificity & moisture requirements

Bryophyte species can be ecologically classified by their substrate dependency (i.e. obligate specialist to generalist) and moisture dependence (i.e. desiccation tolerance, Oliver et al. 2000), which ranges from hydric to mesic to xeric habitats (**Fig. 6**). Over 30 species in GSENM are hydric species requiring aquatic or semi-aquatic habitats with perennial or strong seasonal water availability while mesic species were less diverse, and can survive mild desiccation such as that experienced on rock or soil in high-elevation habitats of GSENM (**Supplement** 4). In contrast, most species in GSENM live commonly in habitats that experience extreme desiccation (xeric), such as soil and rock at low to moderate elevations in shrublands. Many species have moderate substrate specificity being obligate to one or two substrate types, such as calcareous soil and limestone rock, or downed wood and tree bases. Few species are found across multiple substrates except for several “weedy” generalists such as *Syntrichia subpapillosissima* and *Hypnum revolutum* both of which grow on soil, rock, and tree bases (**Fig. 6**).

**Figure 6.**
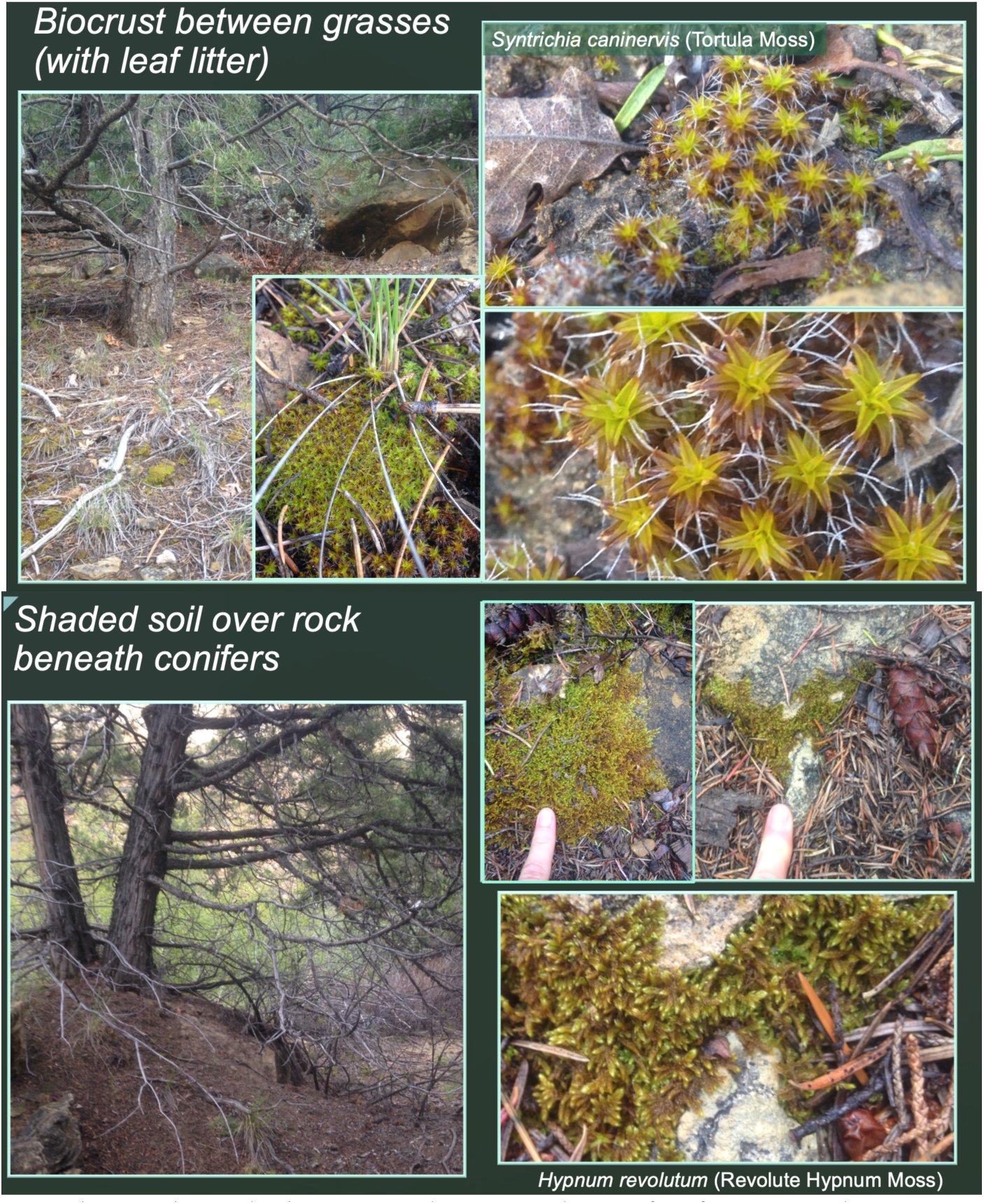
Two mesic bryophyte habitats in the mixed conifer forest with a commonly associated species collected in the Kaiparowits Plateau region in an ephemeral canyon (Site X).

### Site richness patterns & diversity hotspots

Mean site richness was 17.2 ± 9 (SD) and ranged from 4 to 34 species (**Fig. 7**). Six diversity hotspots supported ≥30 species and were canyons with perennial or ephemeral streams having one of four vegetation types (mixed conifer, hardwood-riparian, riparian, or pinyon-juniper).

**Figure 7.**
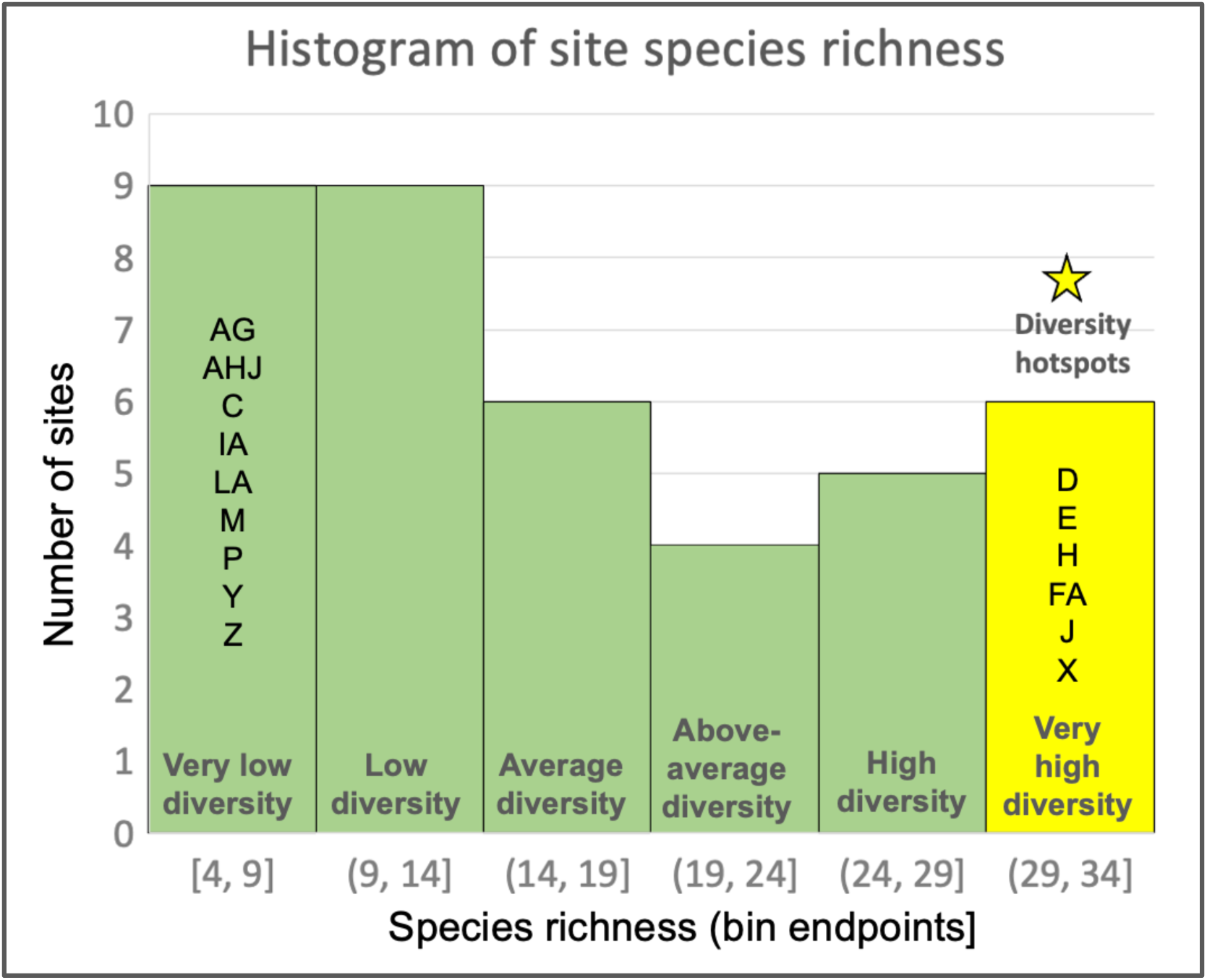
Histogram of estimated species richness across 40 sites (letters listed vertically) in GSENM. Diversity ranged from very low (<9 species) at 9 sites to very high (>29 species) at 6 proclaimed diversity hotspots.

A qualitative ecological assessment of site-level habitat patterns in bryophyte biodiversity is challenging given the broad range of habitat diversity and the low replication within each topo-vegetation combination. Although twenty-five sites were sampled in the Escalante region, 8 in the Kaiparowits Plateau, and 6 in the Staircase (**Table 1**), macrohabitat classes were highly variable in all three regions and likely contribute towards the occurrence of a diversity hotspot in two of the three regions; further collecting to equalize sampling by region will likely reveal hotspots in the Staircase region, in which there are currently none. High richness is likely supported by greater habitat diversity including xeric, mesic, and hydric habitats with variable substrates (e.g. rock, soil, downed wood, and riparian habitat) and unique microhabitats within substrates (e.g. rock crevices and holes). Future analysis will elucidate whether these sites had greater shade (i.e. N-facing canyon walls) and moisture availability (i.e. perennial streams) than other sites, on average.

Diversity hotspots (sites supporting > 25 species) are shown in **Fig. 8** and described in **Table 2** occurred across a broad combination of topography and vegetation classes. This lack of dependency on macrohabitat type suggests that microhabitat diversity may be most important. We suspect that the occurrence of many combinations of substrate and moisture availability within a single site is a feasible explanation for the increased species richness at these localities. Unique combinations of the six topography and seven vegetation classes yielded 19 macrohabitat types across the 40 sites, the most frequently sampled of which were perennial-flowing canyons with riparian or mixed vegetation (deciduous-coniferous riparian) followed by ephemeral-flowing canyons with one of several vegetation types varying in shade and canopy closure (**Table 1**).

**Figure 8.**
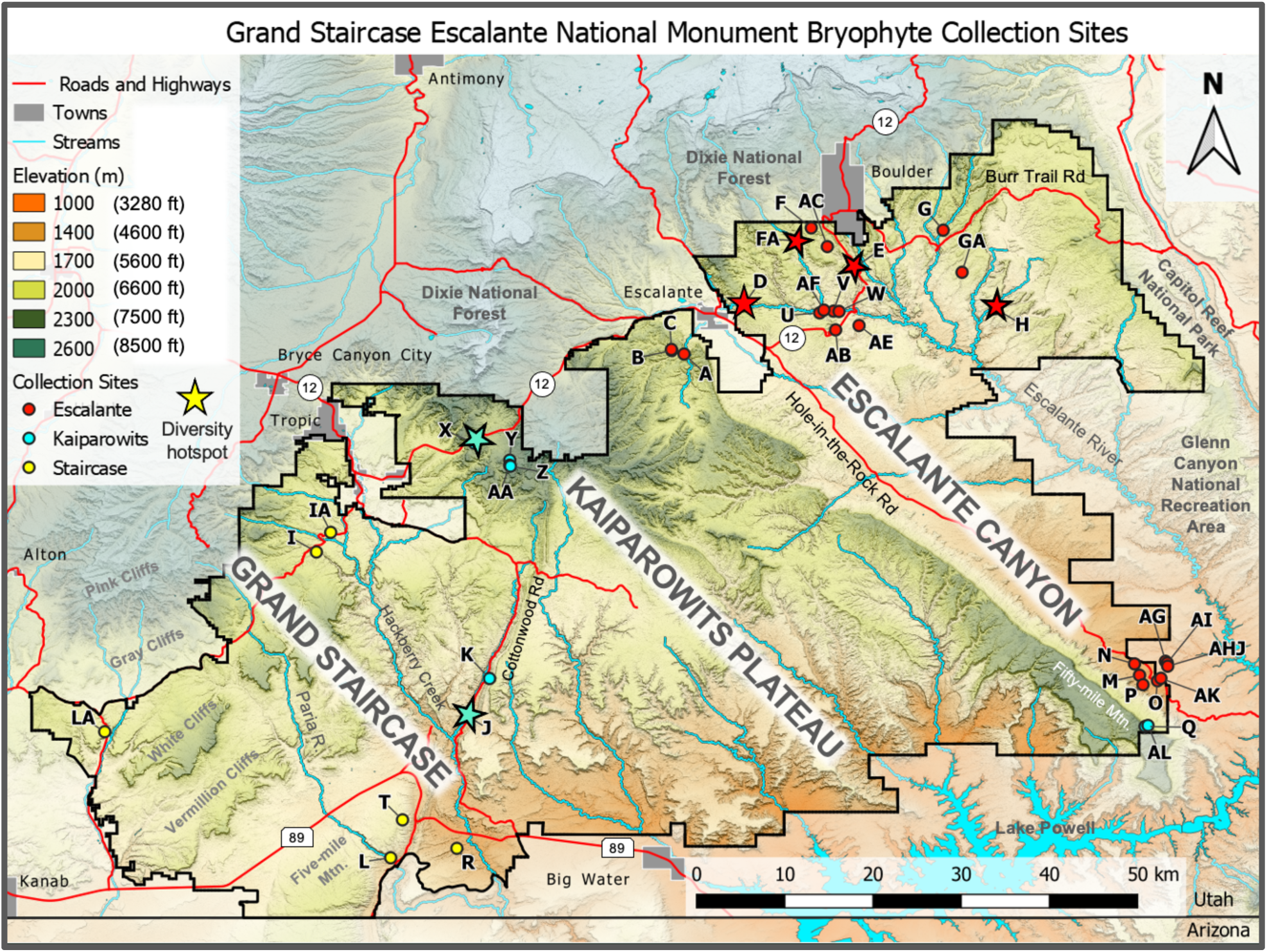
The top six diversity hotspots in GSENM (indicated by stars) included 30 – 34 species in perennial-flowing (**D, E, FA, J**) and ephemeral-flowing (**H, X**) canyons. Notably, these sites are at moderately high elevations >1600 m (5300 ft). Map credit: Tim Wright

### Floristic composition comparison: Grand Canyon vs Grand Staircase

As discussed above, the growth-form and family composition of GSENM bryophytes closely matches that of GCNP despite having 38 fewer total species (**Supplement** 4). At the species level, the two parks share 77 species and of this, 22 are very common, when common is defined as 10 or more collections. **Table 3** lists these 22 shared species all of which can occur on at least three substrate types, but not all of which can be considered generalists (**Fig. 2**). Notably, there are fewer common pleurocarpous mosses than acrocarpous and most of these “feathermosses” are hydric species (**Supplement** 4) common in GSENM only due to the high prevalence of perennial streams, seeps, and springs. In contrast, of the common acrocarpous mosses, only one species is hydric, highlighting a central theme unifying all dryland bryofloras: tree-like acrocarpous mosses are more desiccation tolerant and thus more abundant and frequent than pleurocarpous mosses across mesic and xeric habitats in the Southwest (and elsewhere).

**Table 3.** Common mosses to both GSENM & Grand Canyon National Park, AZ. Common was defined as having a collection frequency ≥10 sites or unique microhabitats.

| Common<br>acrocarpous mosses | Common<br>pleurocarpous mosses |
| --- | --- |
| **Didymodon nevadensis** | **Hypnum revolutum** |
| **\*Didymodon tophaceus** | **Hypnum vaucheri** |
| **Bryoerythrophyllum recurvirostrum** | **\*Conardia compacta** |
| **Tortella alpicola** | **Brachytheciastrum fendleri** |
| **Tortula mucronifolia** | **\*Eucladium verticillatum** |
| **Bryum argenteum** | **\*Eurhynchiastrum pulchellum** |
| **Syntrichia montana** | **Pseudoleskeella tectorum** |
| **Grimmia anodon** |  |
| **Grimmia crinitoleucophaea** |  |
| **Grimmia pulvinata** |  |
| **Grimmia plagiopodia** |  |
| **Encalypta raptocarpa** |  |
| **Syntrichia subpapillosissima** |  |
| **Syntrichia canninervis** | **\*Hydric species** |

However, GCNP has a truly unique *species* composition which comprises 81 taxa not documented for GSENM (**Fig. 9**, **Supplement 4**). Therefore, although total richness in the two parks only differs by 41, the number of species unique to each park is substantial. GCNP has two times the number of unique taxa (**Fig. 9**). Future collecting will inevitably uncover more species for the ecologically rich region of GSENM, however gamma diversity (i.e. landscape-level richness) will likely not reach that of Grand Canyon National Park. The NP harbors ca. 158 species at present, 23% (36 species) of which were found exclusively in the high-elevation mixed conifer forest (Clark 2012). Grand Canyon has more extensive high-elevation habitat. The Kaibab Plateau on the North Rim of the canyon is above 8000 feet (2450 m), while the highest elevation habitat sampled in GSENM was that of Fifty-Mile Mountain (Site Q) and The Blues mountains (Sites AA, Y, and Z) all of which were under 7600 feet (2500 m, **Fig. 10**). Mean richness of the 40 collection sites was < 6000 feet, which often is the upper limit for pinyon-juniper woodlands; this vegetation class has a more open-canopy that increases exposure of bryophytes to solar radiation, and likely limits the establishment of many species at lower elevation (unless there is perennial water source or buffered microhabitats like soil abutting the base of boulders).

**Figure 9.**
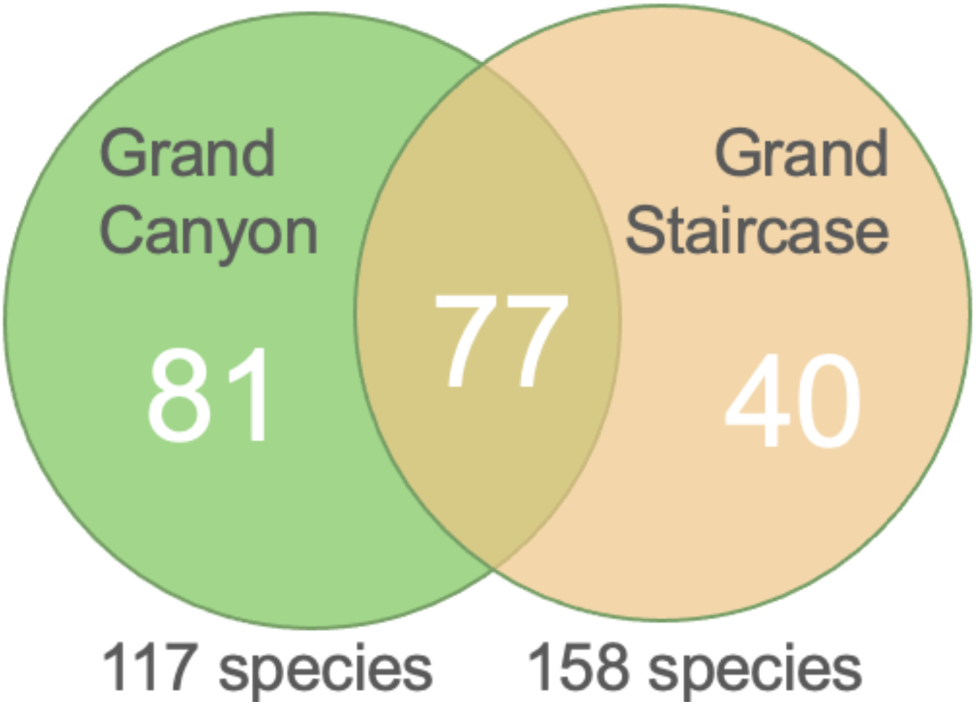
Venn diagram contrasting the number of unique and shared bryophyte species recovered in GCNP (left) and GSENM (right) based on 1500 and 1000 collections, respectively.

**Figure 10.**
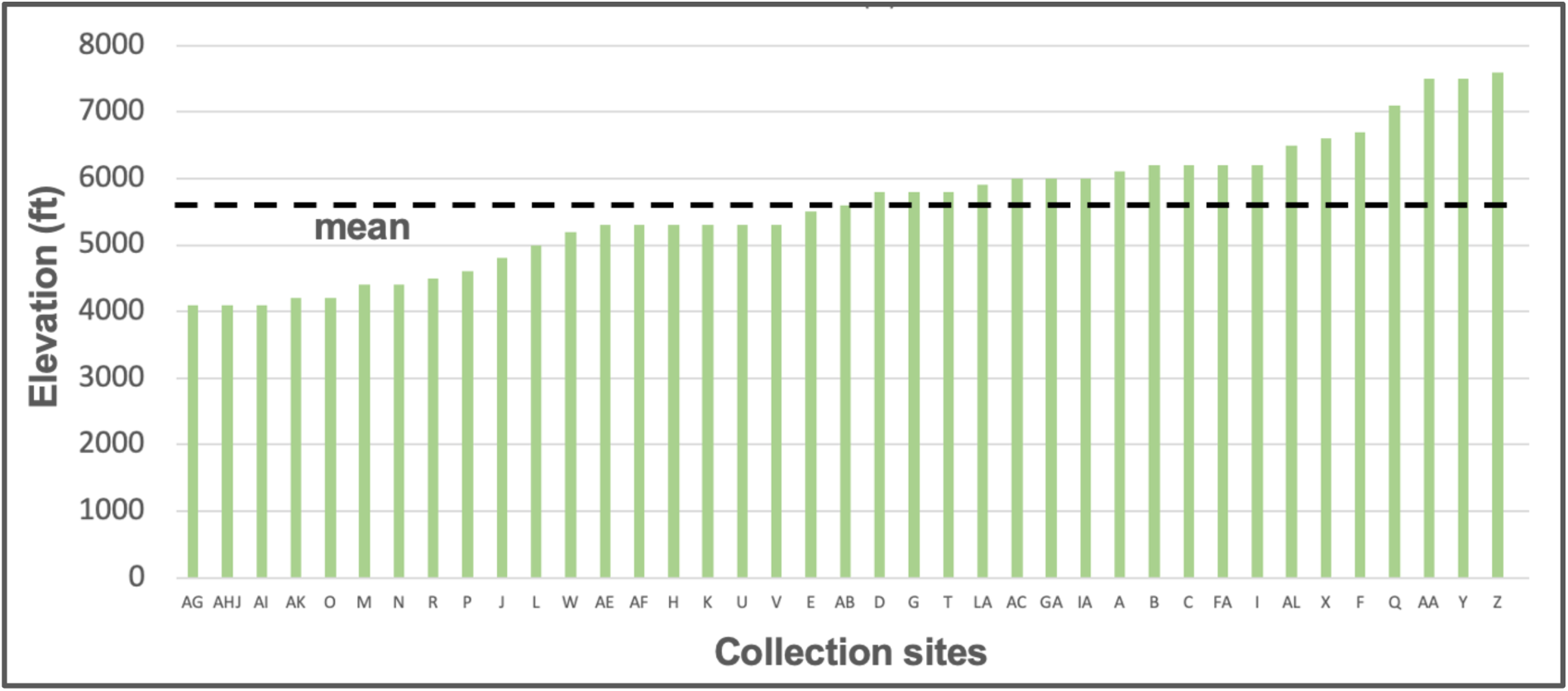
Elevation of bryophyte collection sites in GSENM in increasing order with an overlay of mean site elevation (dashed line).

### Locally rare species and species of conservation concern

In GSENM, ca. 30 species (26%) were found at only one site, while the remaining species were collected between two and 20 times (**Supplement 4**). Most North American bryophytes have broad global distributions spanning multiple continents (BFNA), but this doesn’t exclude the possibility for local rarity. Not surprisingly, rare species often have rare substrates or rare substrate-microclimate combinations (Cleavitt 2005). For example, *Cephaloziella divaricata* is a rare liverwort in drylands in part because of high moisture and shade requirements – the species was found at only one site in GSENM thus far and may be locally rare. Due to the relatively small number of collecting sites in GSENM, however, it may be premature to classify taxa as locally rare at this time. Nonetheless, the current distribution of 17 species in GSENM appears limited in the United States (or globally) and may warrant monitoring in GSENM until further records become available (**Supplement 4**). For example, *Aneura pinguis* (**Fig. 11**) is a common thalloid liverwort in hydric habitats throughout the USA, but its occurrence in GSENM marks a species record for Utah and suggests a need for monitoring to determine potential status of local rarity in Utah.

**Figure 11.**
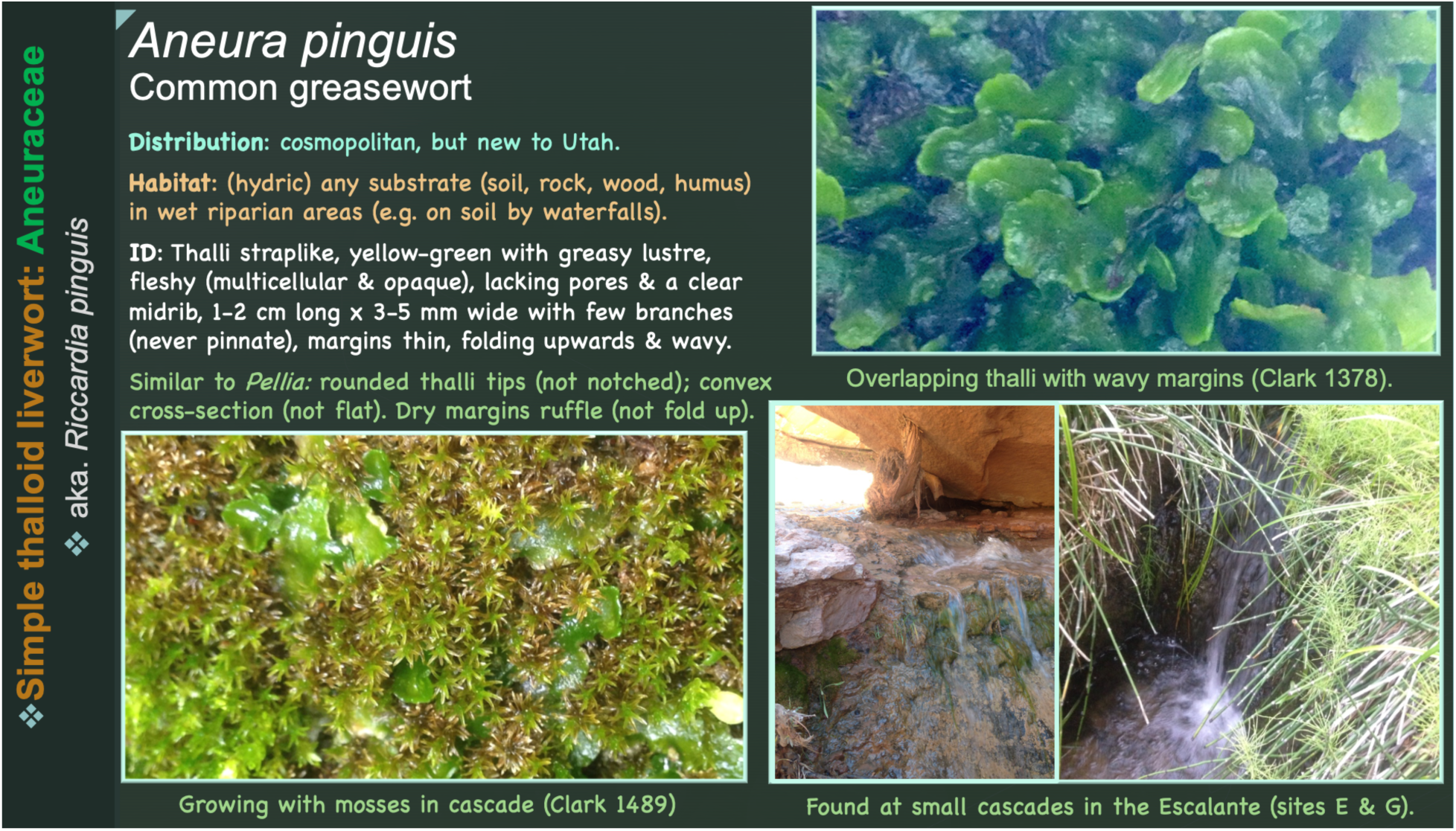
Bryophyte species flashcards like the one above for the common greasewort (*Aneura pinguis*) may help staff and visitors learn to “hunt” for suitable habitat and locally rare thalloid liverwort in GSENM.

## Management recommendations

With potential impacts of climate change and land use on bryophytes in drylands, the following protocol is recommended for GSENM.

### Monitoring bryophytes and diversity hotspots

Due to the strong habitat specificity of bryophytes (e.g. BFNA, Pharo and Beattie 2002), managing and monitoring diversity under future climate change and land-use alterations will necessitate a habitat-stratified approach such as repeated floristic-habitat sampling over time to document changes in site-level richness and to predict other candidate diversity hotspots on the basis of microhabitat-level diversity, which could be assessed by trained non-bryologists. Finding bryophytes in the dried, dormant state (desiccated) requires a careful, trained eye as bryophyte cushions and mats camouflage well on many different substrates (**Fig. 12**).

**Figure 12.**
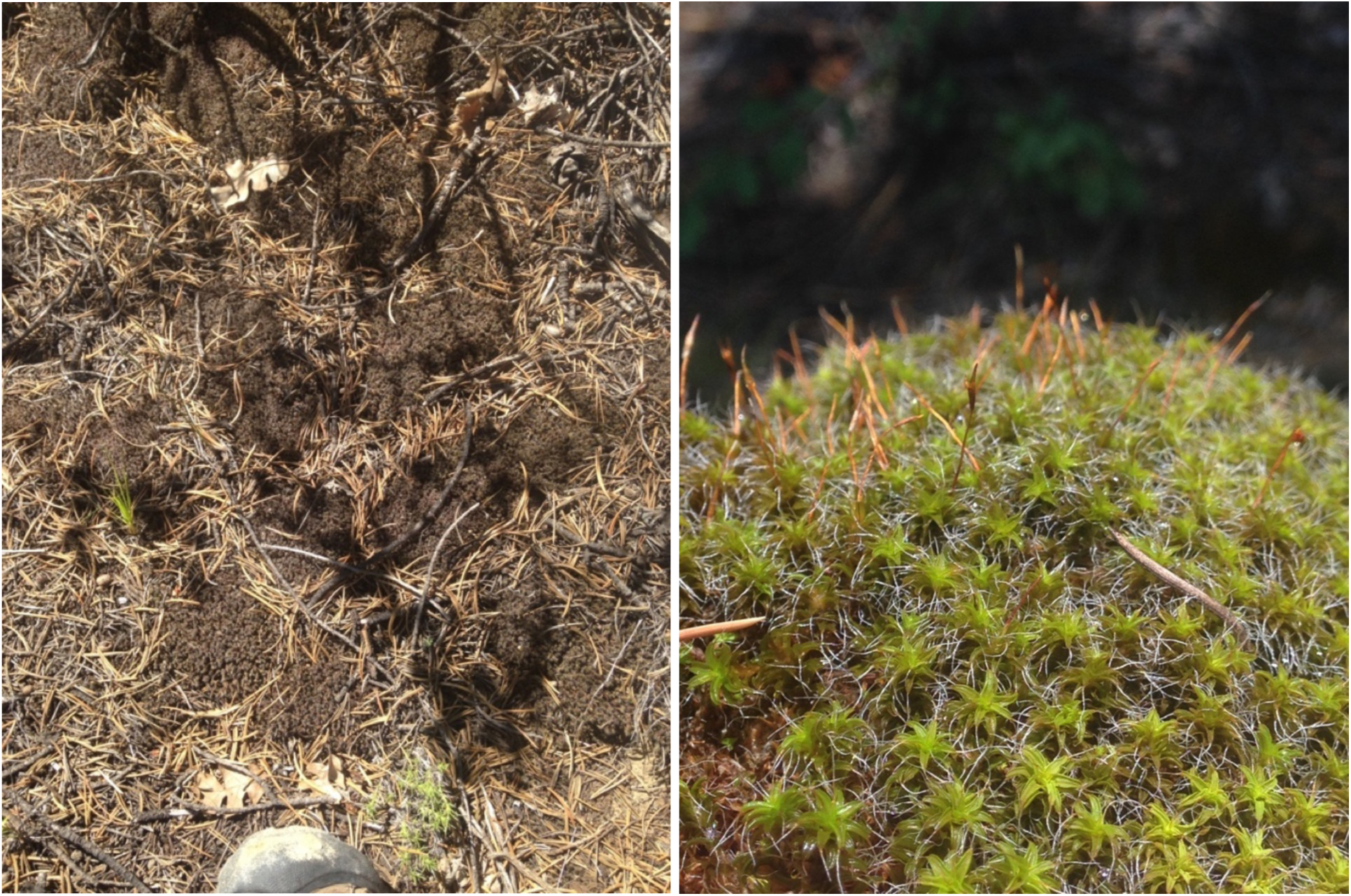
Above a boot (**left**), large cushions of the soil moss, *Syntrichia subpapillosissima*, blend in with pine litter. Wetting small patches (**left**) can aid visual assessment of sexual reproduction (e.g. orange capsules, **right**) and stress, but this should be done conservatively on cloudy days in summer (or preferably not in summer) as short wetting events can cause stress.

### Monitoring biocrust bryophytes

Although biocrusts are not the most diverse microhabitat type for bryophytes at the site-scale, the collective biocrust community across GSENM harbors over 15 species of xeric acrocarpous soil mosses (**Supplement 4**). Many studies recommend monitoring biocrust disturbance from human and animal impact by looking for changes in organismal groups (Bowker et al. 2016), but few have focused on assessing change in species abundance or the photosynthetic health of biocrust mosses (and liverworts in mesic environments). Such assessments are time-intensive and require experts in the field, but may would valuable on a five- or 10-year cycle targeting repeated sampling that would facilitate powerful comparisons over time without the noise introduced by random annual sampling.

Alternatively, a more regular annual monitoring approach could involve a small number of selected microsites to monitor at some of the GSENM hotspot sites (**Fig. 1**) where biocrust species have been found (**Table 2**, **Supplement 3 & 4**). At these hotspots, we recommend targeting a selection of healthy, undisturbed biocrust along with vulnerable microsites, such as those below dead shrubs (**Fig. 13**) or those with fire damage or significant moss canopy deterioration or sun damage (**Fig. 13**). Across landscapes, such disturbed microhabitats are the most likely to show early signs of climate change that could be detected with photographs taken by non-specialists (see instructions on the iNaturalist project, *Citizens of the Crust* (instructions available at https://drylandmoss.wordpress.com/community-outreach/citizen-science/). Photographs could be taken at multiple scales (e.g. 1×1m and 10×10cm) to detect visual signs of stress (**Fig. 13**), compositional shifts in organismal groups, moss growth forms, and even genera or species (**Fig. 14**), if working in tandem with a bryophyte specialist during image analysis.

**Figure 13.**
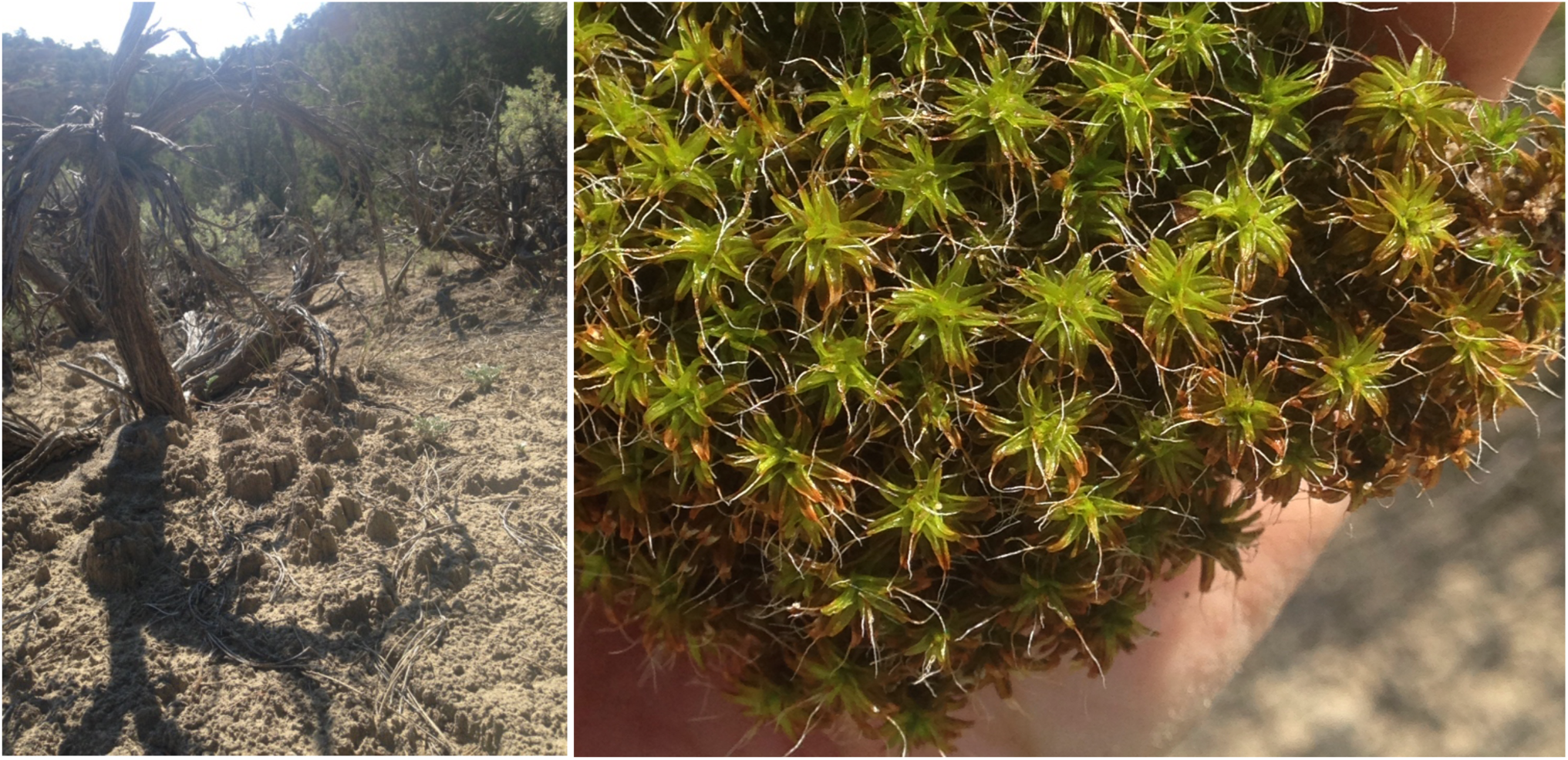
Signs of biocrust moss stress can appear as orange-tipped leaves especially evident when fully hydrated.

**Figure 14.**
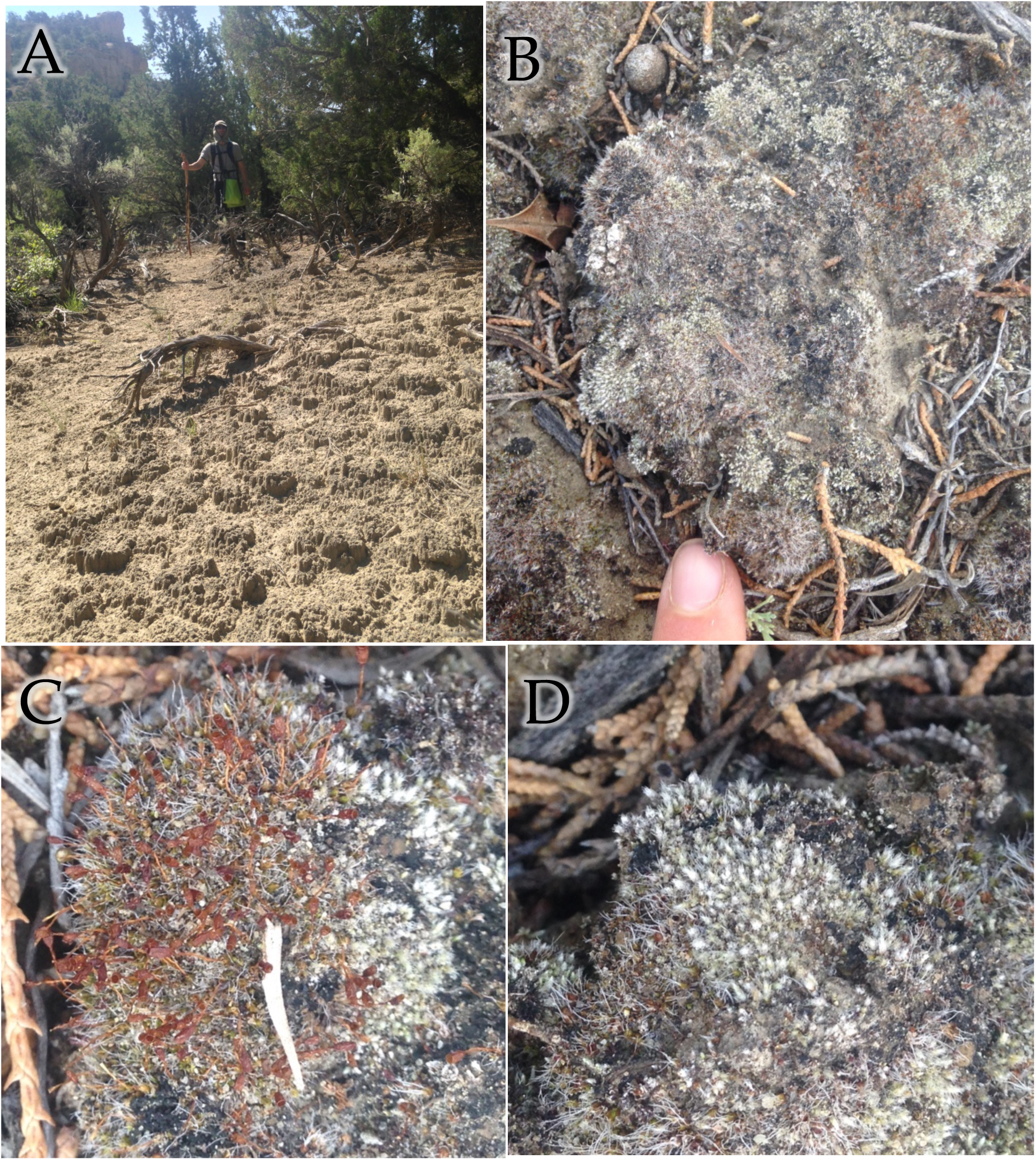
**A.** Finding pinnacled biocrust is the easiest crust type to locate across drylands, but does not support the most diverse biocrust communities. **B.** Low-lying rugose crust often has more species including some of the smallest biocrust acrocarpous species, *Pterygoneurom ovatum* (**C**) and *Bryum lanatum* or *B. argenteum* (**D**).

### Recommendations for GSENM outreach

Increasing visitor awareness about biocrust and dryland bryophytes could involve the following educational outreach elements.

`o` Modern signage for biocrust near trails (macrophotographs)
`o` Bryophyte species flash cards (e.g. **Fig. 10**, https://theresaannclark.wixsite.com/mossesinmotion)
`o` iNaturalist app (e.g. Citizens of the Crust Project & GSENM Project)
`o` Bryophyte interpretative walks (with hand lenses)
`o` Bryophyte/biocrust workshop (with microscopes & local voucher specimens)

### Conclusions & future research

We conclude that the 117 species confirmed for this primary and preliminary bryoflora of Grand Staircase-Escalante National Monument (GSENM) is quite large given relatively few collections (ca. 1000) and a short field investment surveying only 40 sites across ca. 40 days of collecting (**Table 1**). The recently completed vascular plant flora of Grand Staircase-Escalante National Monument (GSENM) also revealed high levels of plant richness exceeding 1100 species (Fertig 2005). Interestingly, the monument appears to support a similar proportion of North American vascular (5.5%) and nonvascular (6.2%) plant species at present (BFNA). This impressive regional plant diversity is partly a consequence of the monument’s location in southern Utah, adjacent to the Mojave and Great Basin Deserts and north of the floristically diverse Kaibab Plateau and Grand Canyon. Another driver of high diversity in GSENM is likely the sheer size of the monument at 1.9 million acres, which increases the probability of supporting more species due to the strong relationship between survey area and species richness.

To ultimately predict bryophyte richness patterns across GSENM, the site data from this project will be used to build landscape and site-scaled models using as predictors GIS climate and habitat data (e.g. soil type, geology, hydrology, shade-time, insolation) and measured environmental data (e.g. vegetation community, microhabitat diversity). Future sampling efforts by the authors or other experts should focus in the Kaiparowits Plateau and Staircase regions where present sampling efforts have been limited. Targeting more north-facing topographical features and perennial sources of water (streams, springs, seeps), which may increase the chances of finding diversity hotspots in these more arid regions of GSENM.

With appropriate support from government institutions like the BLM and a sustained public stewardship, we sincerely hope that the uniquely charismatic and ecologically complex natural lands of GSENM will remain protected thereby allowing the continued study of these important but often overlooked plants into an uncertain future.

## Acknowledgments

The authors graciously acknowledge financial support from the BLM-National Landscape Conservation System (Cooperative Agreement #L14AC00275) along with the guidance of BLM-GSENM Science Program Administrator, Kevin Miller who was especially helpful with many aspects of field logistics. We thank the staff at all visitor centers in Grand Staircase-Escalante National Monument for their assistance with trip planning and amenities that were so critical to field collecting the hot summer. As the primary collector, T. A. Clark is especially grateful for expertise of visiting collector and collaborator, Dr. J. C. Brinda, who helped kickstart a successful summer of collecting in 2015, while also sharing his taxonomic expertise in the field. Field assistants we would like to gratefully acknowledge include Dr. Emma Benenati, Glenn Rink, Emily Warren, and Eric Nolan. We would like to acknowledge Tim Wright for creating the beautiful site map and assistance with acquiring, processing, and summarizing landscape GIS data. Regarding specimen identification, we are indebted to J. C. Brinda for his work at the Missouri Botanical Garden confirming (via microscopic identification) over 50% of species in the present checklist.

---

**Supplement 1.** Catalog of Specimens. This Excel file created by J. C. Brinda contains 395 rows with the following column fields:

|  |  |  |  |  |
| --- | --- | --- | --- | --- |
| Primary collector | Report Sites | Collection number | Collection date | Elev ft |
| Elevation<br>m | Locality | Habitat | Substrate | Associated<br>collectors |
| Taxon name | Determined by | Determination date | Country code | State province |
| County | Management unit |  |  |  |

**Supplement 2.** Catalog of Vouchers. This Excel file created by T. A. Clark contains 800 rows with the following column fields:

|  |  |  |  |  |  |  |
| --- | --- | --- | --- | --- | --- | --- |
| Clark # | GSEM<br>Region | Site | Specimen# | Moss<br>Liverwort | Bryophyte<br>group | Location |
| Family | Species | latitude | longitude | Common<br>name | Additional<br>species | ID notes |
| Elevation<br>m | Habitat | iNaturalist url | Associated<br>Plants | Collectors | Collection<br>Date |  |

**Supplement 3.** Catalog of Sites. This Excel file created by T. A. Clark contains 40 rows with the following column fields:

|  |  |  |  |  |  |  |  |
| --- | --- | --- | --- | --- | --- | --- | --- |
| Region | Site | Locality<br>(details) | Topography | Vegetation | Elev (ft) | Locality description | Date |
| Latitude | Longitude | Collec-<br>tions | Pleuro-<br>carps | Liver-<br>worts | Acrocarps | Total species | Locality |
| # Brinda | # Clark | Soil<br>type | Old Site | Species<br>(code, Sp.) | Bryophyte<br>habitat &<br>community | Site Ecology |  |

**Supplement 4.** Catalog of Species & Traits. This Excel file created by T. A. Clark contains 199 rows with the following column fields:

|  |  |  |  |  |  |  |
| --- | --- | --- | --- | --- | --- | --- |
| GSENM/<br>GCNP | Family | Species | Common<br>synonyms | Growth<br>form | Repro. | Known habitat<br>(adapted from<br>BFNA) |
| Mexico | Canada | Authors | Distribution | Freq<br>GCNP | BFNA or<br>online link | Utah state records |
| Southwest<br>elevation(ft) | Only<br>Hydric | GSENM | GC Parashant | Big Bend<br>NP | Grand<br>Canyon | Southwest endemic |
| Observed<br>habitat) | GSENM<br>collections | Species notes |  |  |  |  |

